# New insights into the mechanisms of plant isotope fractionation from combined analysis of intramolecular ^13^C and deuterium abundances in *Pinus nigra* tree-ring glucose

**DOI:** 10.1101/2024.02.21.581384

**Authors:** Thomas Wieloch, Meisha Holloway-Phillips, Jun Yu, Totte Niittylä

**Author notes:** Corresponding author, Thomas Wieloch. **Author Email address:** Meisha Holloway-Phillips, Jun Yu, Totte Niittylä.

## Abstract

Understanding isotope fractionation mechanisms is fundamental for analyses of plant ecophysiology and paleoclimate based on tree-ring isotope data.

To gain new insights into isotope fractionation, we analysed intramolecular ^13^C discrimination in tree-ring glucose (*Δ_i_*’, *i* = C-1 to C-6) and metabolic deuterium fractionation at H^1^ and H^2^ (*ε*_met_) combinedly. This dual-isotope approach was used for isotope-signal deconvolution.

We found evidence for metabolic processes affecting *Δ*_1_’ and *Δ*_3_’ which respond to air vapour pressure deficit (*VPD*), and processes affecting *Δ*_1_’, *Δ*_2_’, and *ε*_met_ which respond to precipitation but not *VPD*. These relationships exhibit change points dividing a period of homeostasis (1961-1980) from a period of metabolic adjustment (1983-1995). Homeostasis may result from sufficient groundwater availability. Additionally, we found *Δ*_5_’ and *Δ*_6_’ relationships with radiation and temperature which are temporally stable and consistent with previously proposed isotope fractionation mechanisms.

Based on the multitude of climate covariables, intramolecular carbon isotope analysis has a remarkable potential for climate reconstruction. While isotope fractionation beyond leaves is currently considered to be constant, we propose significant parts of the carbon and hydrogen isotope variation in tree-ring glucose originate in stems (precipitation-dependent signals). As basis for follow-up studies, we propose mechanisms introducing *Δ*_1_’, *Δ*_2_’, *Δ*_3_’, and *ε*_met_ variability.

## Introduction

Analysis of the systematic ^13^C/^12^C variation (commonly termed “^13^C signal”; abbreviations in Table 1) across tree-ring series is widely used to study past climate conditions, plant-environment interactions, and physiological traits such as leaf water-use efficiency (CO_2_ uptake relative to H_2_O loss) (Leavitt & Roden, 2022). Signals found at the whole-tissue or whole-molecule level (Fig. 1A, top and middle) are commonly interpreted based on a simplified mechanistic model of ^13^C discrimination, *Δ* (denoting ^13^C/^12^C variation caused by physiological processes) (Farquhar *et al*., 1982). This model considers isotope effects of CO_2_ diffusion from ambient air into intercellular air spaces (Craig, 1953) and CO_2_ assimilation by rubisco (Roeske & O’Leary, 1984) and phospho*enol*pyruvate carboxylase (PEPC; Fig. 2) (Farquhar, 1983; Farquhar & Richards, 1984). Manifestation of these effects as ^13^C discrimination depends on the ratio of intercellular-to-ambient CO_2_ partial pressure (*p*_i_/*p*_a_) (Farquhar *et al*., 1982), and a highly significant positive relationship between *p*_i_/*p*_a_ and leaf *Δ* was confirmed experimentally (Evans *et al*., 1986). Environmental parameters influence *p*_i_/*p*_a_ and thus leaf *Δ* (Evans *et al*., 1986) by affecting the stomatal aperture and CO_2_ assimilation. For instance, in response to drought, isohydric plant species such as *Pinus nigra* (studied here) close their stomata (McDowell *et al*., 2008). This can be expected to decrease *p*_i_/*p*_a_ and leaf *Δ* (Farquhar *et al*., 1982; Evans *et al*., 1986).

**Table 1.** Abbreviations and symbols.

| Abbreviation | Definition |
| --- | --- |
| $^{13}\text{C}$ signal | systematic $^{13}\text{C}/^{12}\text{C}$ variation |
| DAHPS | 3-deoxy-D-arabino-heptulosonate 7-phosphate synthase |
| F6P | fructose 6-phosphate |
| G6P | glucose 6-phosphate |
| G6PD | glucose-6-phosphate dehydrogenase |
| GAP | glyceraldehyde 3-phosphate |
| GAPDH | glyceraldehyde-3-phosphate dehydrogenases |
| NMRS | nuclear magnetic resonance spectroscopy |
| OPPP | oxidative branch of the pentose phosphate pathway |
| PEP | phospho*eno*lpyruvate |
| PEPC | phospho*eno*lpyruvate carboxylase |
| PGA | 3-phosphoglycerate |
| PGI | phosphoglucose isomerase |
| PK | pyruvate kinase |
| RuBP | ribulose 1,5-bisphosphate |
| TCAC | tricarboxylic acid cycle |
| Symbol | Definition |
| $p_a$ | ambient $\text{CO}_2$ partial pressure |
| $p_i$ | intercellular $\text{CO}_2$ partial pressure |
| $PRE$ | precipitation |
| $RAD$ | global radiation |
| $SD$ | sunshine duration |
| $SPEI_i$ | standardised precipitation-evapotranspiration index calculated for different timescales, $i = 1, 3, 6, 8, 12, 16, 24, 36, 48$ months |
| $TMP$ | air temperature |
| $VPD$ | air vapour pressure deficit |
| $\Delta$ | $^{13}\text{C}$ discrimination denoting $^{13}\text{C}/^{12}\text{C}$ variation due to plant physiological processes |
| $\Delta_i'$ | intramolecular $^{13}\text{C}$ discrimination where $i$ denotes individual glucose carbon positions and the prime denotes data corrected for $^{13}\text{C}$ signal redistribution by heterotrophic triose phosphate cycling |
| $\Delta_{1-2}'$ | arithmetic average of $\Delta_1'$ and $\Delta_2'$ |
| $\Delta_{1-3}'$ | arithmetic average of $\Delta_1'$, $\Delta_2'$, and $\Delta_3'$ |
| $\Delta_{5-6}'$ | arithmetic average of $\Delta_5'$ and $\Delta_6'$ |
| $\epsilon_{\text{met}}$ | metabolic deuterium fractionation at glucose $\text{H}^1$ and $\text{H}^2$ |

**Figure 1.**
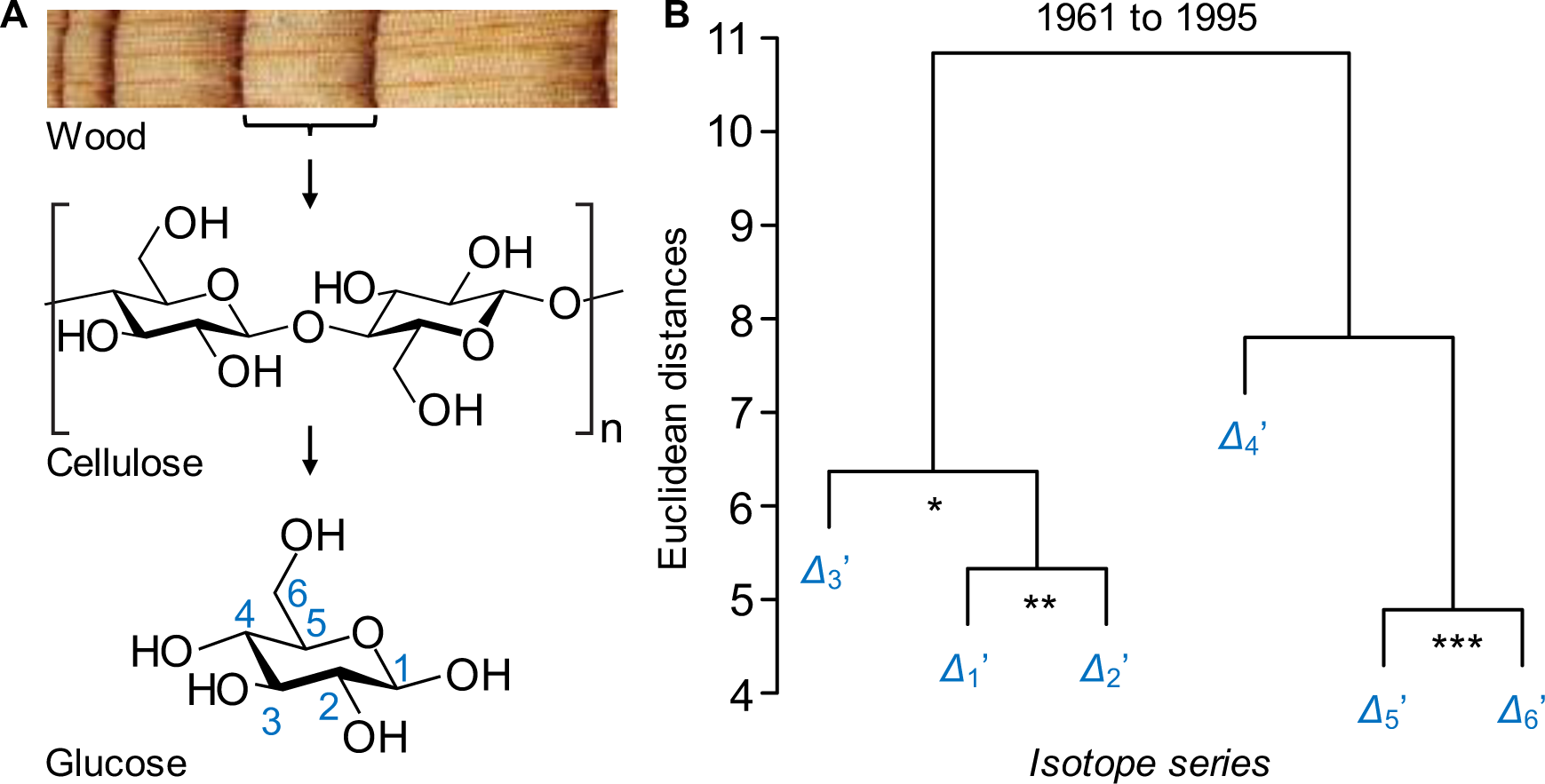
Carbon isotope discrimination in tree rings. **(A)** Levels of resolution of stable carbon isotope analysis: whole plant materials, whole molecules, intramolecular carbon positions. **(B)** Hierarchical clustering of *Δ_i_*’ series for the period 1961 to 1995. Significance of series correlation: *, *p* ≤ 0.05; **, *p* ≤ 0.01; ***, *p* ≤ 0.001. Modified figure from Wieloch *et al*. (2018). *Δ_i_*’ denotes intramolecular ^13^C discrimination in glucose extracted across an annually resolved *Pinus nigra* tree-ring series.

**Figure 2.**
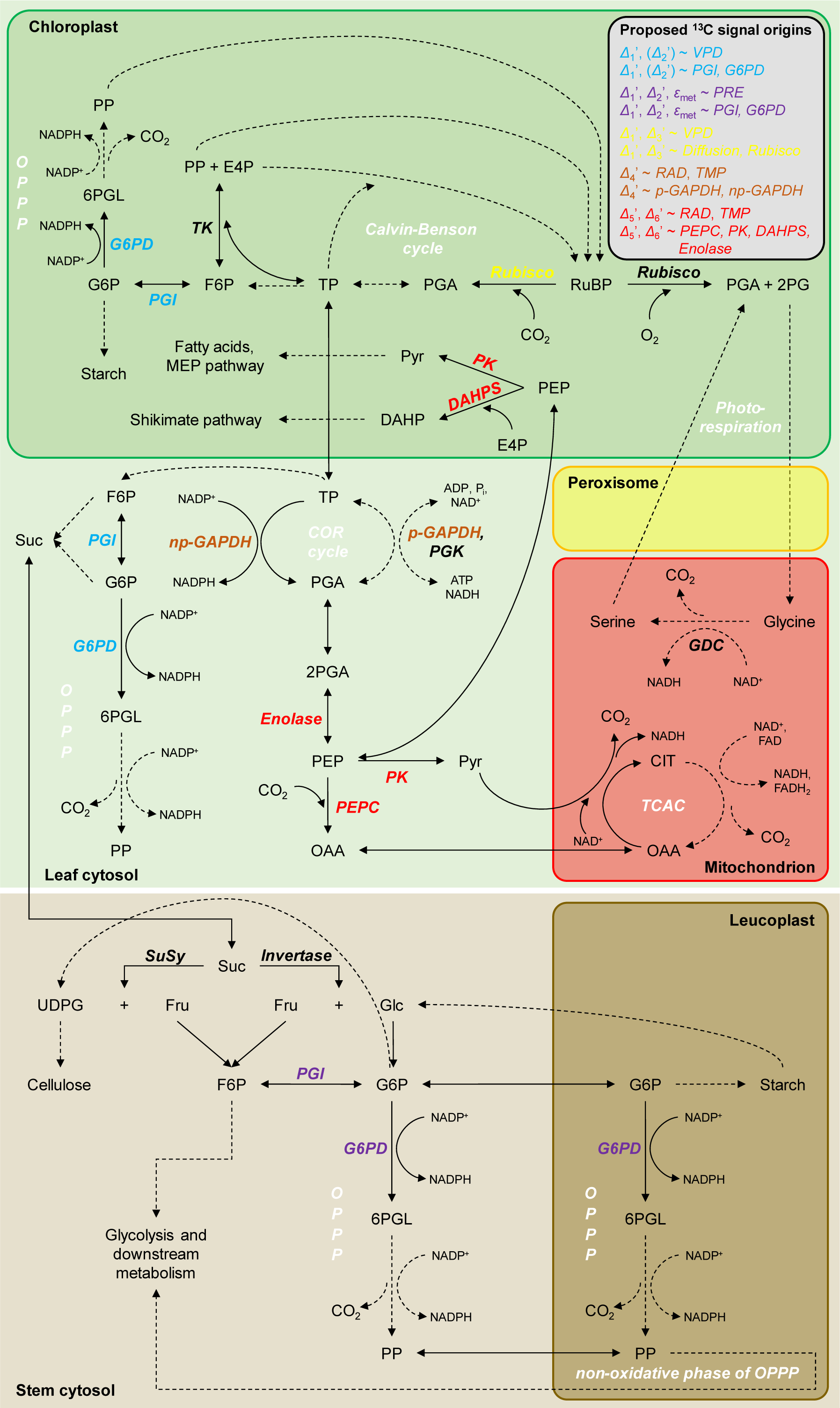
Proposed metabolic origins of carbon and hydrogen isotope signals in tree-ring glucose. Dashed arrows indicate that intermediate reactions are not shown. Abbreviations: 2PG, 2-phosphoglycolate; 2PGA, 2-phosphoglycerate; 6PGL, 6-phosphogluconolactone; ADP, adenosine diphosphate; ATP, adenosine triphosphate; CIT, citrate; COR cycle, cytosolic oxidation-reduction cycle; DAHP, 3-Deoxy-D-*arabino*-heptulosonate 7-phosphate; DAHPS, DAHP synthase; E4P, erythrose 4-phosphate; F6P, fructose 6-phosphate; FAD, flavin adenine dinucleotide; Fru, fructose; G6P, glucose 6-phosphate; G6PD, G6P dehydrogenase; GDC, glycine decarboxylase complex; Glc, glucose; MEP pathway, methylerythritol 4-phosphate pathway; NAD^+^, nicotinamide adenine dinucleotide; NADP^+^, nicotinamide adenine dinucleotide phosphate; np-GAPDH, non-phosphorylating glyceraldehyde-3-phosphate dehydrogenase; OAA, oxaloacetate; OPPP, oxidative pentose phosphate pathway; PEP, phospho*enol*pyruvate; PEPC, PEP carboxylase; p-GAPDH, phosphorylating glyceraldehyde-3-phosphate dehydrogenase; PGA, 3-phosphoglycerate; PGI, phosphoglucose isomerase; PGK, phosphoglycerate kinase; P_i_, inorganic phosphate; PK, pyruvate kinase; PP, pentose phosphate; *PRE*, precipitation; Pyr, pyruvate; Rubisco, ribulose-1,5-bisphosphate carboxylase/oxygenase; RuBP, ribulose 1,5-bisphosphate; *RAD*, global radiation; Suc, sucrose; SuSy, sucrose synthase; TCAC, tricarboxylic acid cycle; TK, transketolase; *TMP*, air temperature; TP, triose phosphates (glyceraldehyde 3-phosphate, dihydroxyacetone phosphate); UDPG, uridine diphosphate glucose; *VPD*, air vapour pressure deficit; *Δ_i_*’, intramolecular ^13^C discrimination where *i* denotes individual glucose carbon positions and the prime denotes data corrected for ^13^C signal redistribution by heterotrophic triose phosphate cycling; *ε*_met_, metabolic deuterium fractionation at glucose H^1^ and H^2^.

Isotope fractionation by metabolic processes downstream of CO_2_ assimilation is complex (Hobbie & Werner, 2004), incompletely understood (Badeck *et al*., 2005; Cernusak *et al*., 2009) and has yet to be adequately integrated into ^13^C-discrimination models (Ubierna *et al*., 2022). Specifically, the simple ^13^C discrimination model described above requires multiple adaptations to enable correct interpretation of the ^13^C composition of tree-ring glucose (studied here). For instance, we recently argued that incorporation of carbon assimilated by PEPC into tree-ring glucose is negligible because leaves lack a high-flux pathway shuttling this carbon into glucose metabolism (Fig. 2) (Wieloch *et al*., 2022c). Therefore, all carbon in tree-ring glucose proposedly derives from rubisco-assimilated CO_2_. Rubisco catalyses the addition of CO_2_ to ribulose 1,5-bisphosphate (RuBP). Since this reaction is essentially the sole carbon source of glucose, ^13^C discrimination accompanying CO_2_ diffusion and subsequent rubisco CO_2_ assimilation (denoted diffusion-rubisco discrimination) is expected to affect all glucose carbon positions equally (Wieloch *et al*., 2018, 2022c).

Moreover, we recently measured *Δ* intramolecularly at all six carbon positions, *i*, of glucose (Fig. 1A, bottom) extracted across an annually resolved tree-ring series of *Pinus nigra* (Wieloch *et al*., 2018). The resultant *Δ_i_*’ dataset comprises 6*31 values (study period: 1961 to 1995; four years missing: 1977, 1978, 1981, 1982) which were corrected for ^13^C signal redistribution by heterotrophic triose phosphate cycling (indicated by prime, Supporting Information Notes S1). We found that, at least, four ^13^C signals contribute to the interannual ^13^C/^12^C variability in tree-ring glucose (Fig. 1B) and proposed the following theories on underlying mechanisms.

We initially proposed the diffusion-rubisco signal is preserved at C-1 to C-3 (Figs. 1B and 2) (Wieloch *et al*., 2018); although this view is modified here. Additionally, C-1 and C-2 are thought to carry ^13^C signals due to fractionation at phosphoglucose isomerase (PGI has carbon isotope effects at both C-1 and C-2) and glucose-6-phosphate dehydrogenase (G6PD has a carbon isotope effect at C-1) (Wieloch *et al*., 2018, 2022a). Two leaf-level mechanisms of signal introduction were proposed. First, with decreasing carbon assimilation, the PGI reaction in chloroplasts moves from being on the side of fructose 6-phosphate (F6P) towards equilibrium (Fig. 2) (Dietz, 1985). This shift is expected to cause ^13^C enrichments at C-1 and C-2 of glucose 6-phosphate (G6P) and its derivatives starch and tree-ring glucose (Table 2) (Wieloch *et al*., 2018). Moreover, shifts towards PGI equilibrium are associated with G6P increases (Dietz, 1985). Increasing G6P is thought to cause G6PD activation and thus increasing flux through the oxidative pentose phosphate pathway (OPPP) in chloroplasts (Cossar *et al*., 1984; Sharkey & Weise, 2016; Preiser *et al*., 2019) resulting in additional ^13^C enrichment at C-1 of G6P and its derivatives (Wieloch *et al*., 2022a). Hydrogen isotope evidence consistent with these proposed metabolic shifts was reported recently (Wieloch *et al*., 2022a). Second, the PGI reaction in chloroplasts is usually displaced from equilibrium on the side of F6P whereas the PGI reaction in the cytosol is closer to or in equilibrium (Dietz, 1985; Gerhardt *et al*., 1987; Leidreiter *et al*., 1995; Schleucher *et al*., 1999; Szecowka *et al*., 2013). This is expected to result in ^13^C/^12^C differences between starch and sucrose at both hexose C-1 and C-2 (Table 2) (Wieloch *et al*., 2022b). By extension, changes in the relative contribution of starch to the biosynthesis of tree-ring glucose is expected to contribute to the ^13^C signals at C-1 and C-2.

**Table 2.**
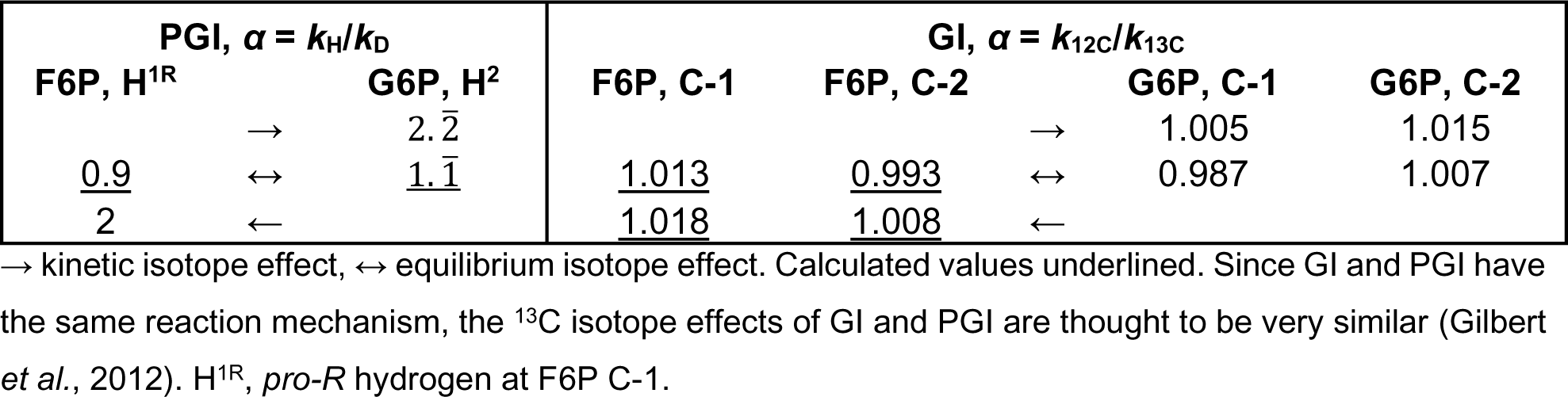
Hydrogen isotope effects of phosphoglucose isomerase (PGI, Rose & O’Connell, 1961), and carbon isotope effects of glucose isomerase (GI, Gilbert *et al*., 2012).

In addition to ^13^C signals at C-1 and C-2, tree-ring glucose samples discussed here carry deuterium signals caused by metabolic processes at H^1^ and H^2^. These signals are strongly correlated and were approximated as

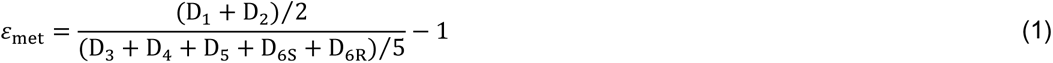

where D*_i_* denotes relative deuterium abundances at individual H-C positions (Wieloch *et al*., 2022a,b). Variability of *ε*_met_ pertaining to glucose H^1^ and H^2^ was attributed to isotope effects of G6PD (*k*_H_⁄*k*_D_ = 2.97) (Hermes *et al*., 1982) and PGI (Table 2) (Rose & O’Connell, 1961; Wieloch *et al*., 2022a,b), respectively. Proposedly, G6PD and PGI-dependent metabolic processes in both leaves and tree rings may contribute to *ε*_met_ signal introduction (Wieloch *et al*., 2022b). Interestingly, Wacker (2022) recently reported that the commonly observed whole-molecule deuterium depletion of leaf starch which derives from deuterium depletion at starch glucose H^2^ (Schleucher *et al*., 1999; Wieloch *et al*., 2022a) is not detectable in nocturnal sucrose. Proposedly, this depletion is either washed out at the level of cytosolic PGI or masked either by the vacuolar sucrose pool or deuterium enrichments at other sucrose hydrogen positions. Washout would imply that any *ε*_met_ signal present at leaf-level G6P H^2^ is lost to the medium. In this case, the *ε*_met_ signal at tree-ring glucose H^2^ may originate outside of leaves.

At tree-ring glucose C-4 (Fig. 1B), the diffusion-rubisco ^13^C signal is thought to be absent due to counteracting fractionation by leaf-cytosolic glyceraldehyde-3-phosphate dehydrogenases (GAPDH; Fig. 2) (Wieloch *et al*., 2021). Signal removal may involve both changes in 3-phosphoglycerate (PGA) flux into downstream metabolism including the tricarboxylic acid cycle (TCAC) relative to flux into tree-ring glucose and changes of flux through the cytosolic oxidation-reduction cycle (Wieloch, 2021; Wieloch *et al*., 2021).

The ^13^C signal at C-5 and C-6 (Fig. 1B) is thought to derive from the postulated (but not yet measured) isotope effects of leaf-level enzymes that modify the carbon double bond in phospho*enol*pyruvate (PEP, Fig. 2) (Wieloch *et al*., 2022c). This includes enolase, pyruvate kinase (PK), PEPC, and 3-deoxy-D-*arabino*-heptulosonate-7-phosphate synthase (DAHPS), the first enzyme of the shikimate pathway. Breaking the double bond in PEP is thought to proceed faster when ^12^C instead of ^13^C forms this bond (Wieloch *et al*., 2022c). Consequently, increasing relative flux into metabolism downstream of PEP is thought to ^13^C enrich remaining PEP at the double-bond carbons and their derivatives including glucose C-5 and C-6 (Wieloch *et al*., 2022c). For example, O_3_ causes downregulation of rubisco, upregulation of PEPC, and DAHPS expression (Dizengremel, 2001; Janzik *et al*., 2005; Betz *et al*., 2009). This is expected to cause increasing relative flux into metabolism downstream of PEP (Wieloch *et al*., 2022c). Accordingly, we previously found a negative relationship between reconstructed tropospheric O_3_ concentration and tree-ring glucose *Δ*_5-6_’ (arithmetic average of *Δ*_5_’ and *Δ*_6_’, Table 1) (Wieloch *et al*., 2022c).

By contrast, the diffusion-rubisco signal is not evident at C-5 and C-6 (Wieloch *et al*., 2022c). This was explained (*inter alia*) by interaction between photorespiration and the TCAC (Fig. 2, Wieloch *et al*., 2022c). Photorespiration increases with drought which results in increasing supply of mitochondrial NADH via the glycine decarboxylase complex. Since this NADH can feed oxidative phosphorylation, NADH and FADH_2_ supply by the TCAC which requires injection of PEP into the TCAC via PK and PEPC may be reduced. This should result in *Δ*_5-6_’ increases counteracting drought-induced decreases in diffusion-rubisco discrimination (see above).

The theories of isotope signal introduction outlined above require further testing. They derive from separate analyses of either the *Δ_i_*’ or deuterium dataset. However, some reactions exhibit both carbon and hydrogen isotope effects (e.g., G6PD at G6P C-1 and H^1^; PGI at G6P C-1, C-2, and H^2^ but not H^1^) and should therefore introduce intercorrelated ^13^C and deuterium signals (suggested terminology: hydro-carbon isotope signals and hydro-carbon isotope fractionation). Combined analysis of intramolecular ^13^C and deuterium data can, in principle, help to separate those signals from signals introduced by reactions which merely exhibit either carbon or hydrogen isotope effects. Therefore, we here studied the relationships between *Δ_i_*’ and *ε*_met_ and their dependence on environmental parameters. Based on our results, we critically examine and revise existing isotope theory and provide new insights into a central open question—whether carbon and hydrogen isotope variability across tree rings derives from leaf-level processes only (as supported by current evidence) or whether processes in the stem contribute as well.

## Material and Methods

The *Δ_i_*’ and *ε*_met_ datasets reanalysed here are described in Wieloch *et al*. (2018, 2022b) and in Notes S1. Data of relative humidity, precipitation (*PRE*), global radiation (*RAD*), sunshine duration (*SD*), and air temperature (*TMP*) are from the climate station Hohe Warte (Vienna, Austria, 48.23° N, 16.35° E, 198 m AMSL) (Klein Tank *et al*., 2002). Air vapour pressure deficit (*VPD*) was calculated following published procedures (Abtew & Melesse, 2013). Data of the standardised precipitation-evapotranspiration index (*SPEI_i_*) calculated for integrated periods of *i* = 1, 3, 6, 8, 12, 16, 24, 36, 48 months were obtained for 48.25° N, 16.25° E (Fan & van den Dool, 2004; Beguería *et al*., 2010). The *SPEI* is a multi-scalar drought index approximating soil moisture variability when calculated for short timescales and groundwater variability when calculated for long timescales (Vicente-Serrano *et al*., 2010). The *RAD* series starts in 1964 while all other climate series start in 1961. Horizontal distances between the tree site and the climate station and grid point are < 15 km. Vertical offsets are small. Hence, climate data and site conditions are expected to be in good agreement. Analytical procedures are described in Notes S1.

## Results

### Hydro-carbon isotope signals at tree-ring glucose HC-1 and HC-2

Tree-ring glucose of our *Pinus nigra* samples exhibits strongly correlated hydrogen isotope signals at H^1^ and H^2^ (Wieloch *et al*., 2022b). These signals occur only after crossing a change point in 1980. Isotope-environment-relationship analyses indicated that the trees had likely access to groundwater before 1980 which prevented changes of the processes introducing these isotope signals. We proposed the signals derive from the hydrogen isotope effects of G6PD (*k*_H_⁄*k*_D_ = 2.97) (Hermes *et al*., 1982) and PGI (Table 2) (Rose & O’Connell, 1961; Wieloch *et al*., 2022a) in autotrophic and/or heterotrophic tissue (Fig. 2; ‘Introduction’) (Wieloch *et al*., 2022a,b). If this proposal is correct then there should be related signals in *Δ*_1_’ and *Δ*_2_’ due to the carbon isotope effects of G6PD affecting C-1 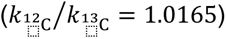 (Hermes *et al*., 1982) and PGI affecting C-1 and C-2 (Table 2) (Gilbert *et al*., 2012). Several findings support this hypothesis. First, among all *Δ_i_*’ series, *Δ*_1_’, *Δ*_1-2_’, *Δ*_1-3_’, and *Δ* are not normally distributed (Table S1, negative skew). Second, among these nonnormal series, *Δ*_1-2_’, *Δ*_1-3_’, and *Δ* exhibit a change point in 1980 (*Δ*_1-2_’: parametric test, *p* < 0.001, nonparametric test, *p* < 0.05; *Δ*_1-3_’: parametric test, *p* < 0.01; *Δ*: parametric test: *p* < 0.05; *n* = 31). Third, 1983 to 1995 average values of *Δ*_1_’, *Δ*_2_’, *Δ*_1-2_’, and *Δ* are significantly lower than the average values of 1961 to 1980, while the 1983 to 1995 variance is significantly larger (Table S2). By contrast, *Δ*_3_’ does not exhibit significant differences in average value or variance between the two periods. Fourth, *Δ*_1_’ and *Δ*_2_’ data pertaining to 1983 to 1995 are significantly correlated (*r* = 0.67, *p* = 0.01, *n* = 13). Fifth, *ε*_met_ approximates average hydrogen isotope fractionation at glucose H^1^ and H^2^ caused by metabolic processes (Eq. 1). Using simple linear regression modelling, we found significant negative relationships between the 1983 to 1995 data of *ε*_met_ and *Δ*_1_’ as well as *Δ*_2_’, but not *Δ*_3_’ or any other *Δ_i_*’ (Fig. 3, green circles; *Δ*_1_’ ∼ *ε*_met_: *R^2^* = 0.35, *adjR^2^* = 0.29, *p* = 0.03; *Δ*_2_’ ∼ *ε*_met_: *R^2^* = 0.54, *adjR^2^* = 0.50, *p* = 0.004; *Δ*_3_’ ∼ *ε*_met_: *R^2^* = 0.21, *adjR^2^* = 0.13, *p* > 0.1; *n* = 13; Table S3). Our ^13^C-NMRS data exhibit relatively large measurement errors. Based on estimates of this random error variance, about 88% of the variance in the *Δ*_1_’ and *Δ*_2_’ data of 1983 to 1995 is systematic variance (Table S4). Hence, about 33% and 57% of the systematic variance in *Δ*_1_’ and *Δ*_2_’ is explained by processes causing *ε*_met_ variation (0.29/0.88 and 0.5/0.88) while about 67% and 43%, respectively, go back to other processes. Taken together, carbon and hydrogen isotope signals at glucose HC-1 and HC-2 are significantly associated during 1983 to 1995 but not during 1961 to 1980 (Notes S2). The processes introducing these signals cause concerted ^13^C and deuterium enrichments (Fig. 3A-B).

**Figure 3.**
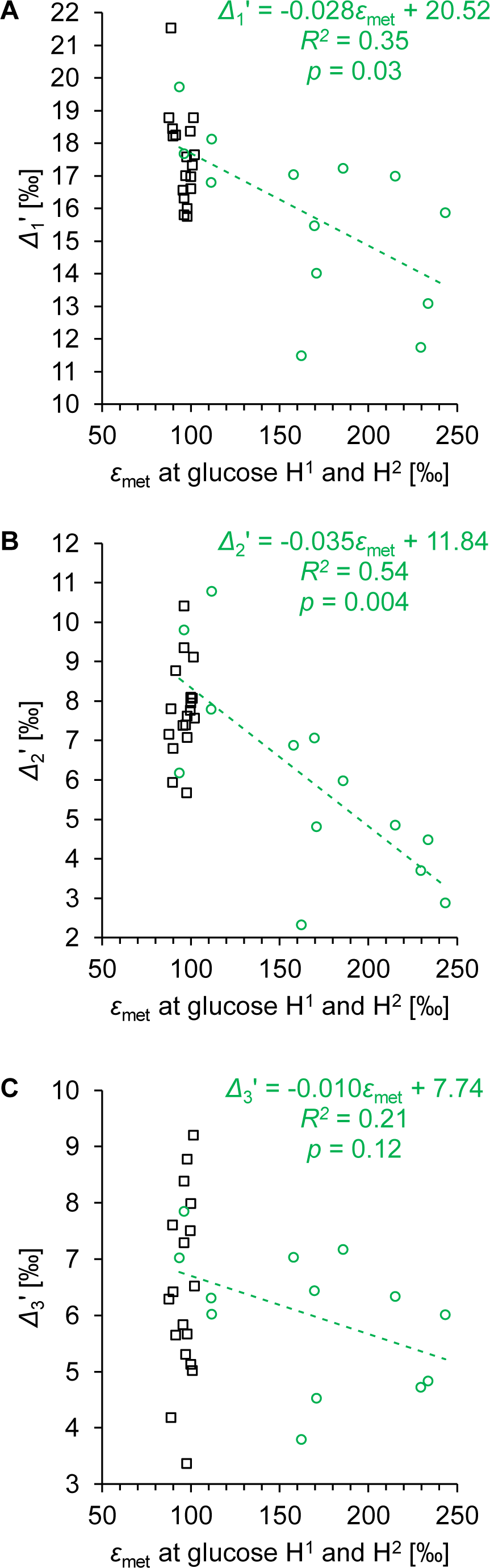
Relationship between the average hydrogen isotope fractionation caused by metabolic processes at glucose H^1^ and H^2^ (*ε*_met_) and ^13^C discrimination at C-1, C-2, and C-3 (*Δ*_1_’, *Δ*_2_’, and *Δ*_3_’). Glucose was extracted across an annually resolved tree-ring series of *Pinus nigra* from the Vienna Basin (black squares, 1961 to 1980; green circles, 1983 to 1995). Dashed line, relationship between the hydrogen and carbon isotope data of the period 1983 to 1995.

### Isotope-environment relationships at tree-ring glucose C-1 to C-3

As evident from our previously published hierarchical cluster analysis and Pearson correlation analyses for the whole period (1961 to 1995), *Δ*_1_’, *Δ*_2_’, and *Δ*_3_’ share common variability (Fig. 1B) (Wieloch *et al*., 2018). Since *Δ*_1-2_’ and *Δ*_1-3_’ exhibit change points in 1980 (see above and Tables S1-2), we analysed the early (1961 to 1980) and late period (1983 to 1995) separately.

During the late period, *Δ*_1_’ and *Δ*_3_’ are more closely associated (Fig. 4A; *r* = 0.87, *p* = 10^−4^, *n* = 13) than *Δ*_1_’ and *Δ*_2_’ (*r* = 0.67, *p* = 0.01, *n* = 13). While this contrasts with results for the whole period (Fig. 1B), it is consistent with isotope-climate-relationship patterns for the late period. *Δ*_1_’ and *Δ*_3_’ correlate similarly with numerous climate parameters and periods (Table 3; *VPD*, *PRE*, *SPEI*_1_ to *SPEI*_16_, *TMP*, *SD*). By contrast, *Δ*_2_’ correlates only with one *VPD* period and several *PRE* periods. A model including *ε*_met_ and growing season *VPD* as cofactors captures most of the systematic variance in *Δ*_1_’ of 88% (Table 4, M1; Table S4). Consistent with the findings above (Fig. 3, Table 3), only *ε*_met_ but not growing season *VPD* contributes significantly to the *Δ*_2_’ model whereas only growing season *VPD* but not *ε*_met_ contributes significantly to the *Δ*_3_’ model (Table 4, M2-3). Removing insignificant terms, we find that *ε*_met_ explains 57% of the systematic variance in *Δ*_2_’, while growing season *VPD* explains the entire systematic variance in *Δ*_3_’ (Table 4, M4-5; Table S4). The effect of *VPD* on *Δ*_1_’ is about twice as large as on *Δ*_3_’ (Table 4, M1 versus M5) while the effect of *ε*_met_ on *Δ*_1_’ is about half as large as on *Δ*_2_’ (M1 versus M4). Intriguingly, *Δ*_1_’ and *Δ*_3_’ are affected by processes that respond to growing season *VPD*. *VPD*-dependent processes can account for both the clustering and correlation between *Δ*_1_’ and *Δ*_3_’ data of 1983 to 1995 (Fig. 4A). By contrast, *ε*_met_ is significantly correlated only with *PRE* (especially March to July *PRE*) but no other climate parameter (Table S5; Table 4, M11). Furthermore, in our *Δ*_1_’ and *Δ*_2_’ models, *ε*_met_ can be substituted by March to July *PRE* (Table 4, M1 versus M6, M4 versus M7).

**Figure 4.**
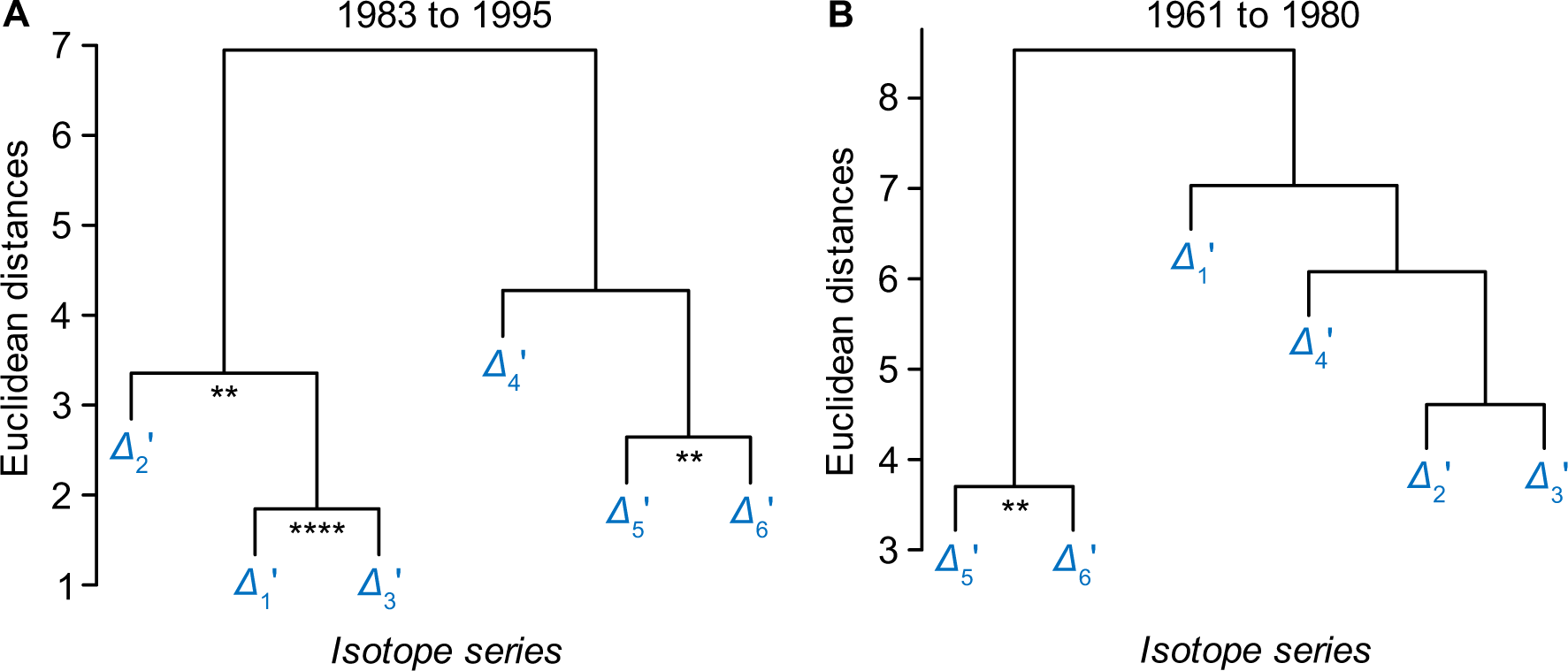
Hierarchical clustering of *Δ_i_*’ series for the periods 1983 to 1995 (A) and 1961 to 1980 (B). *Δ_i_*’ denotes intramolecular ^13^C discrimination in tree-ring glucose of *Pinus nigra* from the Vienna basin with *i* denoting individual glucose carbon positions. Significance of series correlation: **, *p* ≤ 0.01; ****, *p* ≤ 10^−4^.

**Table 3.**
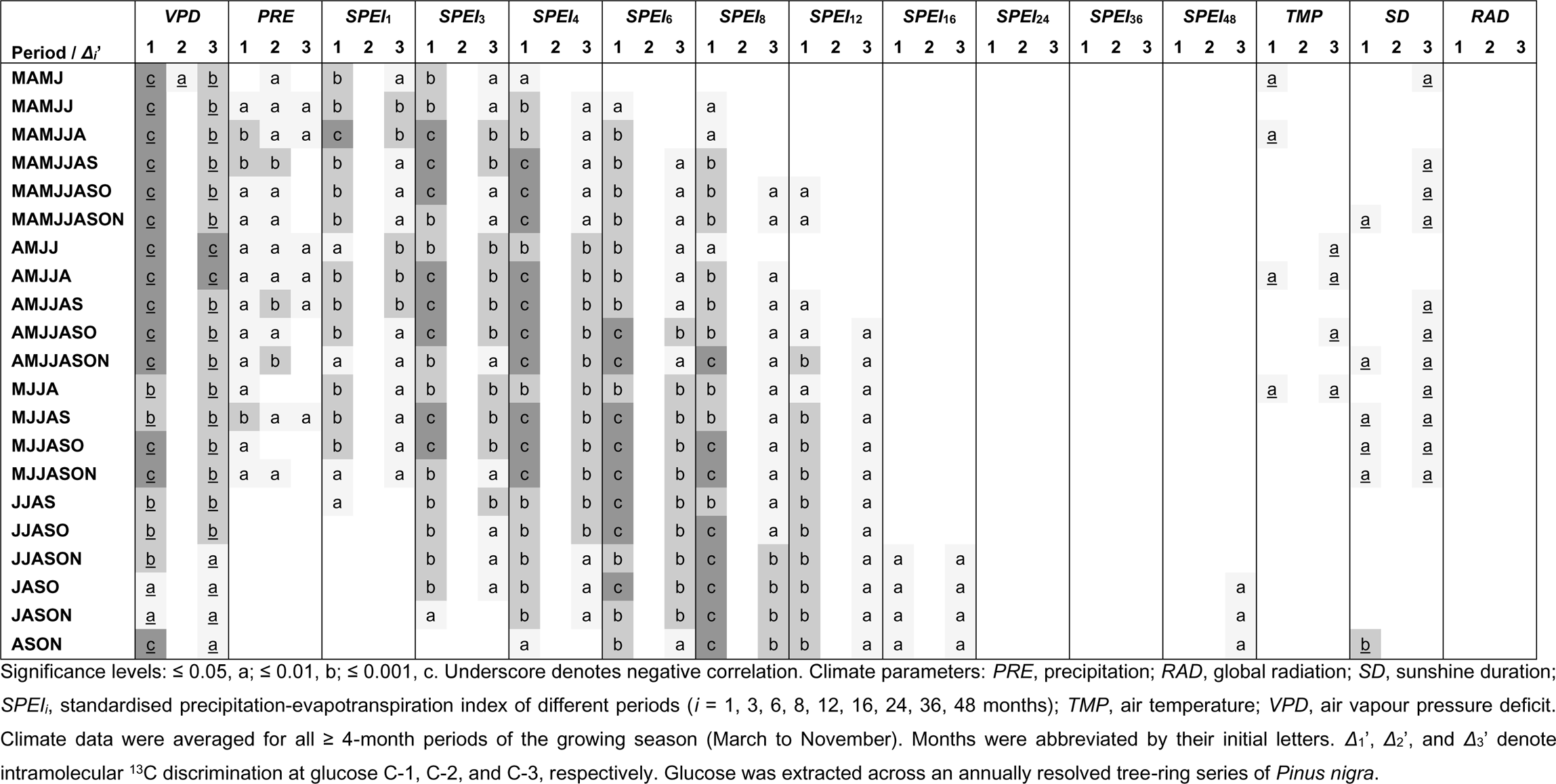
Significance of Pearson correlations among *Δ*_1_’, *Δ*_2_’, and *Δ*_3_’ and climate series for the period 1983 to 1995 (*n* = 13).

**Table 4.**
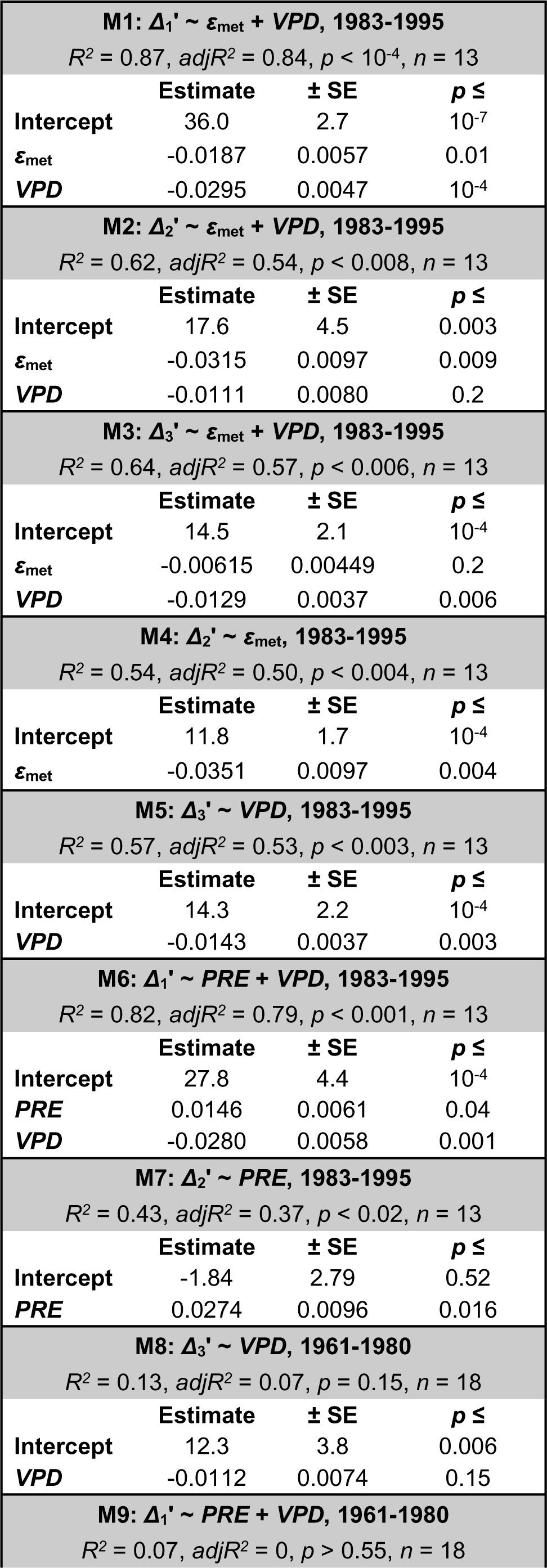
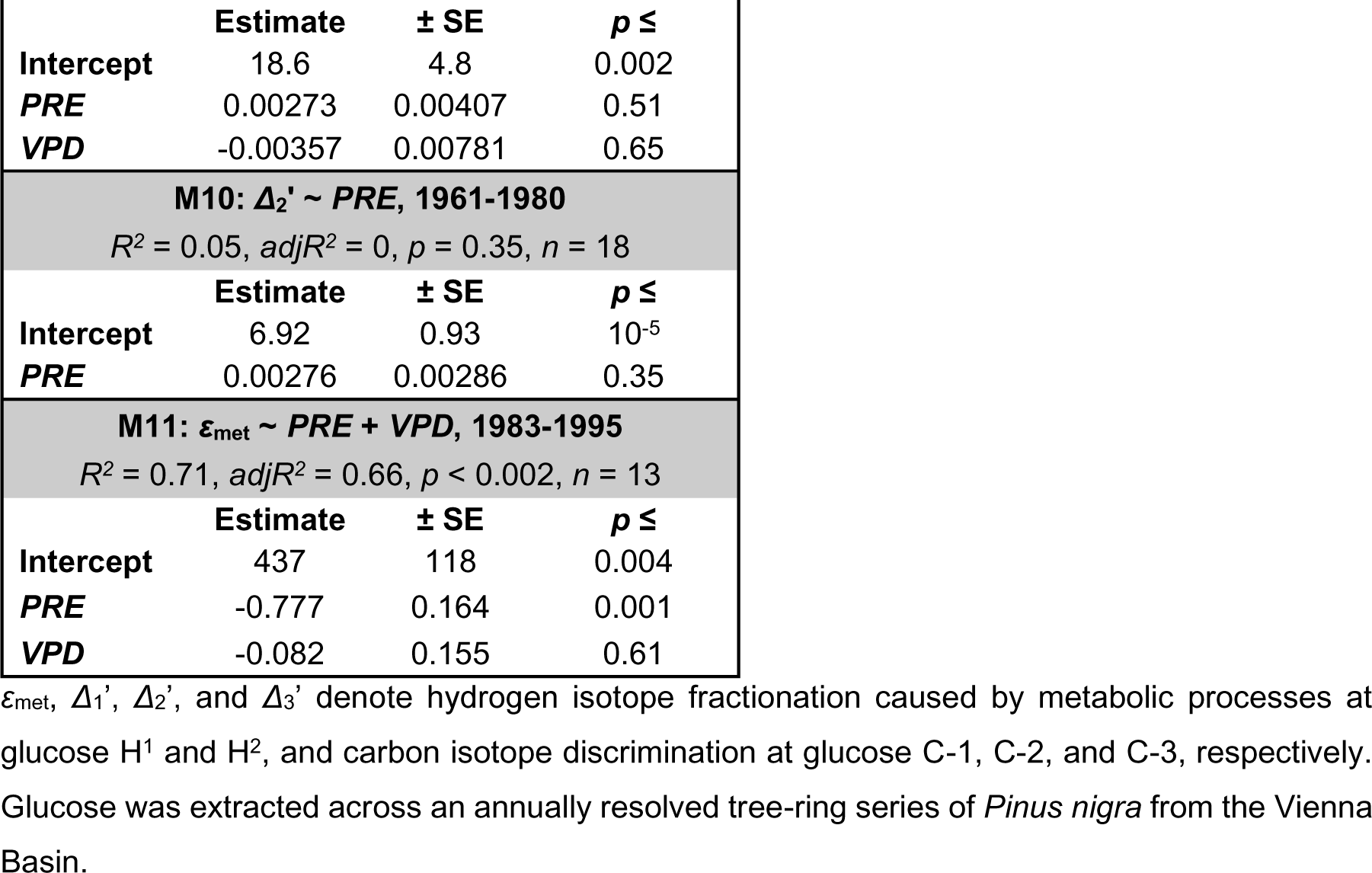
Linear regression models of *Δ*_1_’, *Δ*_2_’, *Δ*_3_’, and *ε*_met_ as function of *ε*_met_, growing season air vapour pressure deficit (*VPD*) and March to July precipitation (*PRE*).

During the early period, *Δ*_1_’, *Δ*_2_’, and *Δ*_3_’ are not significantly correlated (Fig. 4B). Furthermore, isotope-environment models that work for the late period (Table 4, M5-7) do not work for the early period (M8-10). Compared to the late period, we found fewer and weaker isotope-climate correlations (Table 5).

**Table 5.**
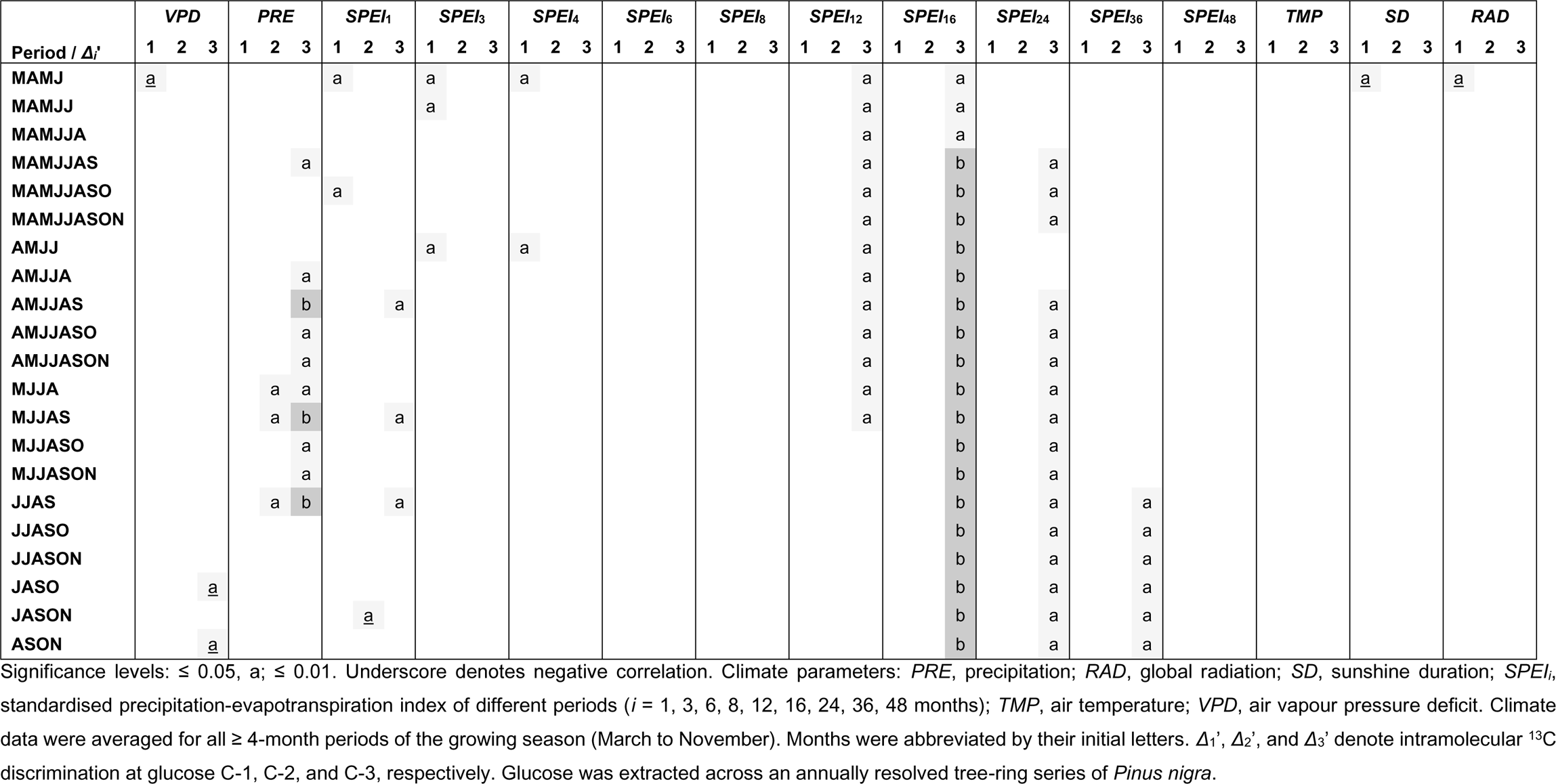
Significance of Pearson correlations among *Δ*_1_’, *Δ*_2_’, and *Δ*_3_’ and climate series for the period 1961 to 1980 (*n* = 18).

### Isotope-environment relationships at tree-ring glucose C-4 to C-6

As evident from our previously published hierarchical cluster analysis and Pearson correlation analyses for the whole period, *Δ*_4_’, *Δ*_5_’, and *Δ*_6_’ share common variability, and *Δ*_5_’ and *Δ*_6_’ are significantly correlated (Fig. 1B; *r* = 0.61, *p* < 0.001, *n* = 31) (Wieloch *et al*., 2018). This significant correlation holds for both the early and late period (Fig. 4). Furthermore, we did not find change points in the *Δ*_4_’, *Δ*_5_’, and *Δ*_6_’ series (see also Tables S1-2). Therefore, we analysed isotope-environment relationships for the whole period. We found that *Δ*_5_’ and *Δ*_6_’ correlate with numerous climate parameters and periods but most significantly with *RAD* while significant *Δ*_4_’-climate correlations are rare (Table 6). Models including April to September *RAD* and March to October *TMP* as cofactors capture 96% of the systematic variance in *Δ*_5-6_’, *Δ*_5_’, and *Δ*_6_’ of 73%, 66%, and 45%, respectively (Table 7, M1-3; *Δ*_5-6_’, *adjR^2^*= 0.70, *p* = 10^−7^; *Δ*_5_’, *adjR^2^* = 0.64, *p* = 10^−6^; *Δ*_6_’, *adjR^2^* = 0.43, *p* < 0.001; *n* = 28; Table S4). Based on *RAD* regression slopes (which are better constrained than *TMP* regression slopes), the ^13^C discrimination at C-5 is about 1.5 times larger than at C-6 (Table 7, M2-3). The model works well for both the early and late period (Table 7, M4-5). Furthermore, consistent with the weak association between *Δ*_4_’ and *Δ*_5-6_’ (Fig. 1B), the model works reasonably well for *Δ*_4_’ considering the relatively low systematic variance in *Δ*_4_’ of 38% (Table 7, M6; Table S4).

**Table 6.**
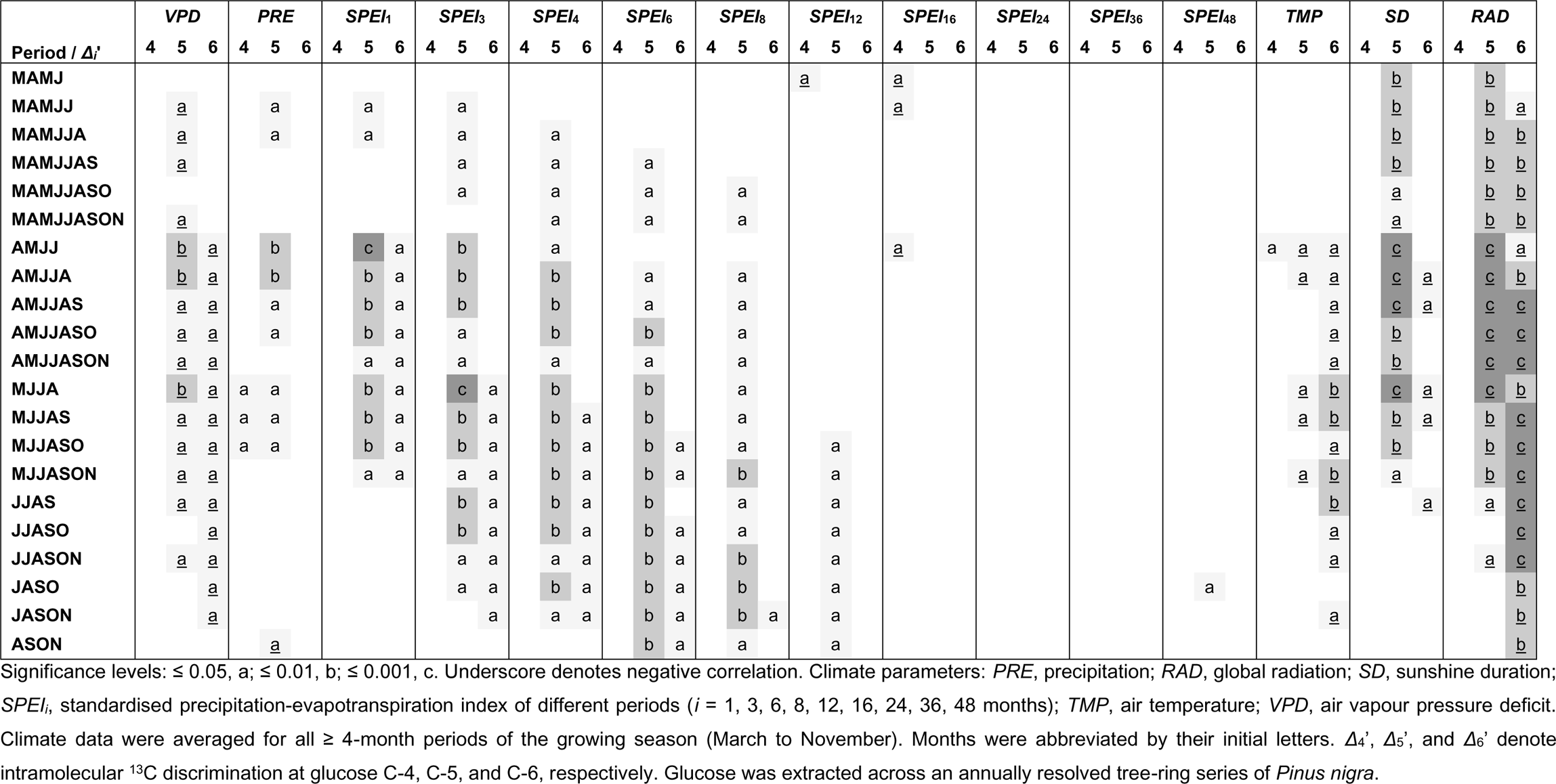
Significance of Pearson correlations among *Δ*_4_’, *Δ*_5_’, and *Δ*_6_’ and climate series for the period 1961 to 1995 (*n* = 31).

**Table 7.** Linear regression models of *Δ*_4_’, *Δ*_5_’, and *Δ*_6_’ as function of April to September global radiation (*RAD*), and March to October air temperature (*TMP*).

| **M1: $\Delta_{5-6}' \sim RAD + TMP$, 1964-1995** |  |  |  |
| --- | --- | --- | --- |
| $R^2 = 0.72$, $adjR^2 = 0.70$, $p = 10^{-7}$, $n = 28$ | | | |
|  | **Estimate** | **± SE** | **$p \leq$** |
| **Intercept** | 26.0 | 3.1 | $10^{-8}$ |
| ***RAD*** | -0.00843 | 0.00105 | $10^{-7}$ |
| ***TMP*** | 1.35 | 0.29 | $10^{-4}$ |
| **M2: $\Delta_5' \sim RAD + TMP$, 1964-1995** |  |  |  |
| $R^2 = 0.66$, $adjR^2 = 0.64$, $p = 10^{-6}$, $n = 28$ | | | |
|  | **Estimate** | **± SE** | **$p \leq$** |
| **Intercept** | 24.8 | 4.3 | $10^{-5}$ |
| ***RAD*** | -0.0103 | 0.0015 | $10^{-6}$ |
| ***TMP*** | 1.81 | 0.40 | $10^{-4}$ |
| **M3: $\Delta_6' \sim RAD + TMP$, 1964-1995** |  |  |  |
| $R^2 = 0.47$, $adjR^2 = 0.43$, $p < 0.001$, $n = 28$ | | | |
|  | **Estimate** | **± SE** | **$p \leq$** |
| **Intercept** | 27.3 | 4.2 | $10^{-6}$ |
| ***RAD*** | -0.00658 | 0.00144 | $10^{-4}$ |
| ***TMP*** | 0.876 | 0.393 | 0.04 |
| **M4: $\Delta_{5-6}' \sim RAD + TMP$, 1964-1980** |  |  |  |
| $R^2 = 0.69$, $adjR^2 = 0.63$, $p < 0.001$, $n = 15$ | | | |
|  | **Estimate** | **± SE** | **$p \leq$** |
| **Intercept** | 31.2 | 6.7 | 0.001 |
| ***RAD*** | -0.00906 | 0.00177 | 0.001 |
| ***TMP*** | 1.11 | 0.47 | 0.04 |
| **M5: $\Delta_{5-6}' \sim RAD + TMP$, 1983-1995** |  |  |  |
| $R^2 = 0.82$, $adjR^2 = 0.79$, $p < 0.001$, $n = 13$ | | | |
|  | **Estimate** | **± SE** | **$p \leq$** |
| **Intercept** | 29.1 | 4.5 | $10^{-4}$ |
| ***RAD*** | -0.00875 | 0.00132 | $10^{-4}$ |
| ***TMP*** | 1.22 | 0.40 | 0.01 |
| **M6: $\Delta_4' \sim RAD + TMP$, 1964-1995** |  |  |  |
| $R^2 = 0.15$, $adjR^2 = 0.09$, $p = 0.12$, $n = 28$ | | | |
|  | **Estimate** | **± SE** | **$p \leq$** |
| **Intercept** | 8.33 | 4.73 | 0.09 |
| ***RAD*** | -0.00266 | 0.00160 | 0.11 |
| ***TMP*** | 0.931 | 0.439 | 0.04 |

$\Delta_4'$ , $\Delta_5'$ , and $\Delta_6'$ denote carbon isotope discrimination at glucose C-4, C-5, and C-6, respectively. Glucose was extracted across an annually resolved tree-ring series of *Pinus nigra* from the Vienna Basin.

## Discussion

### Intramolecular carbon isotope analysis of tree-ring glucose yields information about metabolic variability and water status of both leaves and stems

We found evidence for processes affecting *Δ*_1_’ and *Δ*_3_’ which respond to *VPD* (Table 4, M1, M3, M5). Intriguingly, we also found evidence for processes simultaneously affecting *ε*_met_, *Δ*_1_’, and *Δ*_2_’ which respond to *PRE* but not *VPD* (Table S5; Table 4, M1, M2, M4, M6, M7, M11). This sensitivity to different hydrological properties may be explained by the fact that stem capacitance can buffer stem water status against changes in *VPD* (McCulloh *et al*., 2019), whereas leaf water status is tightly coupled to *VPD* (Grossiord *et al*., 2020). Changes in *PRE* will affect soil water potential and hence both stem and leaf water status. Variability in leaf water status may be impacted more by *VPD* than by soil water status which would explain why *VPD* is the best predictor of the intercorrelated processes affecting *Δ*_1_’ and *Δ*_3_’. By contrast, *VPD*-insensitive processes affecting *ε*_met_, *Δ*_1_’, and *Δ*_2_’ may reside in stems. Hence, we propose intramolecular carbon and hydrogen isotope analysis of tree-ring glucose yields information about metabolic variability and water status not only of leaves but also of stems. *PRE*-dependent systemic changes in enzyme expression can be considered as an alternative explanation.

### Isotope fractionation mechanisms in leaves affecting tree-ring glucose C-1 to C-3

The *Δ*_1-2_’ and *Δ*_1-3_’ series exhibit change points in 1980, i.e., their frequency distributions do not align with the properties of a single theoretical probability distribution (‘Results’; Tables S1-2). Consequently, we investigated the early (1961 to 1980) and late period (1983 to 1995) separately. During the late period, *Δ*_1_’ and *Δ*_3_’ are significantly intercorrelated (Fig. 4A) and correlate negatively with *VPD* and positively with short-term *SPEI* whereas *Δ*_2_’ lacks most of these correlations (Table 3). Furthermore, during the late period, growing season *VPD* accounts for a significant fraction of the systematic variance in *Δ*_1_’ and the entire systematic variance in *Δ*_3_’ but does not contribute significantly to explaining *Δ*_2_’ (Table 4, M1, M2, M5; Table S4). Hence, increasing *VPD* during 1983 to 1995 causes ^13^C enrichments at tree-ring glucose C-1 and C-3 but not C-2. At C-1, the effect is about twice as large as at C-3 (Table 4, M1 and M5).

As discussed above, the *VPD*-dependent processes affecting *Δ*_1_’ and *Δ*_3_’ are likely located in leaves. Qualitatively, *VPD*-induced ^13^C enrichments at C-1 and C-3 are consistent with the mechanisms of diffusion-rubisco fractionation (see ‘Introduction’). However, diffusion-rubisco fractionation affects all glucose carbon positions equally (Wieloch *et al*., 2018). Hence, the unequal *VPD* response of *Δ*_1_’, *Δ*_2_’, and *Δ*_3_’ points to post-rubisco fractionations. In the following, we assume *Δ*_3_’ variation derives entirely from diffusion-rubisco fractionation and argue *VPD*-dependent isotope fractionation at PGI and G6PD in leaf chloroplasts and the cytosol may exert additional control over *Δ*_1_’ and *Δ*_2_’ variability. Generally, variability in PGI fractionation depends on three biochemical properties: (i) the equilibrium status of the PGI reaction, and relative flux of the PGI reactants (ii) F6P and (iii) G6P into competing metabolic pathways (Figs. 2 and 5):

(i) PGI reversibly converts F6P into G6P (Fig. 5A). Under nonstress conditions, the PGI reaction in chloroplasts is strongly displaced from equilibrium on the side of F6P (Dietz, 1985; Gerhardt *et al*., 1987; Kruckeberg *et al*., 1989; Schleucher *et al*., 1999; Wieloch *et al*., 2022a; Wieloch, 2022). With decreasing *p*_i_, however, the reaction moves towards equilibrium (Dietz, 1985; Wieloch *et al*., 2022a). This shift is accompanied by ^13^C increases at C-1 and C-2 of G6P (Table 2) which will be transmitted to downstream derivatives such as starch and tree-ring glucose (Wieloch *et al*., 2018). In isohydric species such as *Pinus nigra*, *p*_i_ decreases with drought due to stomatal closure (McDowell *et al*., 2008). Here, we found stronger *VPD*-induced ^13^C increases at tree-ring glucose C-1 than at C-3. This is consistent with the PGI-related isotope shift expected at C-1. However, the apparent absence of the diffusion-rubisco signal from C-2 contrasts with the expected isotope shift. That said, in *Phaseolus vulgaris*, the ratio of leaf sucrose-to-starch carbon partitioning was shown to increase steeply with decreasing *p*_i_ (Sharkey *et al*., 1985). Hence, the relative contribution of chloroplastic G6P and its isotope composition to downstream metabolism may decrease with increasing *VPD* reducing the influence of the mechanism described on *Δ*_1_’ and *Δ*_2_’ variation.
(ii) In natural systems, leaf nighttime respiration is increased under drought (Fig. 5B; Schmiege *et al*., 2023). Furthermore, in the dark, the cytosolic PGI reaction was found to be near equilibrium (Gerhardt *et al*., 1987). Consequently, F6P would be ^13^C depleted at C-1 but ^13^C enriched at C-2 relative to the corresponding G6P positions (Table 2). Increasing relative F6P flux into mitochondrial respiration would then result in ^13^C increases at C-1 and ^13^C decreases at C-2 of G6P and downstream derivatives. Thus, this mechanism is consistent with both observations, stronger *VPD*-induced ^13^C increases at tree-ring glucose C-1 compared to C-3 and the apparent absence of the diffusion-rubisco signal from C-2.
(iii{) While carbon assimilation commonly decreases with drought (McDowell *et al*., 2008), the activity of leaf-cytosolic G6PD increases (Fig. 5C; Landi *et al*., 2016). This can be expected to result in increasing relative G6P flux into the OPPP. While some authors reported that the cytosolic PGI reaction in illuminated leaves is in equilibrium (Gerhardt *et al*., 1987) others found displacements from equilibrium (Leidreiter *et al*., 1995; Schleucher *et al*., 1999; Szecowka *et al*., 2013). Hence, PGI-related isotope shifts in tree-ring glucose resulting from G6P flux into the leaf-cytosolic OPPP are hard to predict (Table 2). By contrast, the unidirectional conversion of G6P to 6-phosphogluconolactone catalysed by G6PD proceeds faster with ^12^C-1 than ^13^C-1 G6P 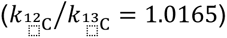 (Hermes *et al*., 1982). Hence, increasing relative flux through the leaf-cytosolic OPPP may contribute to the stronger *VPD*-induced ^13^C increases at tree-ring glucose C-1 compared to C-3.

**Figure 5.**
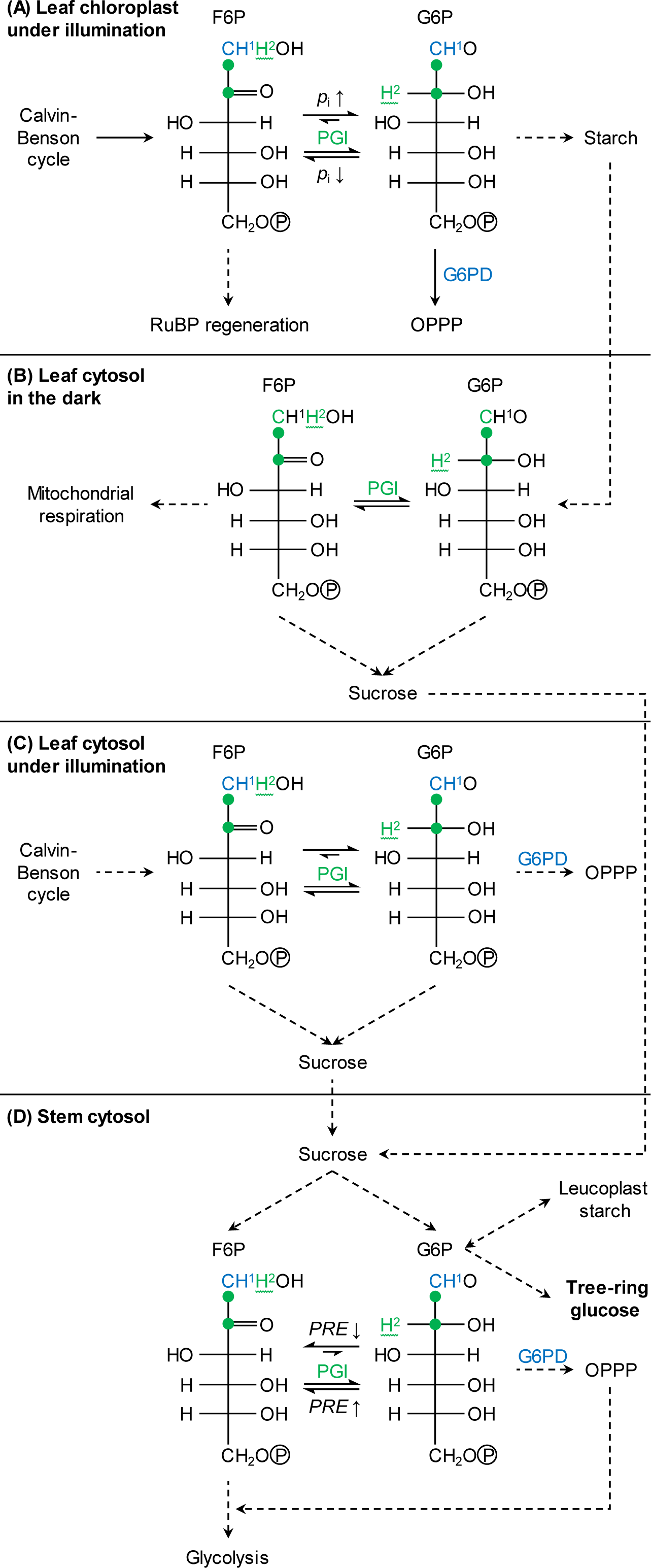
Processes invoked to explain isotope fractionation at tree-ring glucose HC-1 and HC-2: (A) in leaf chloroplasts under illumination, (B) in the leaf cytosol in the dark, (C) in the leaf cytosol under illumination, and (D) in the stem cytosol. F6P and G6P carbon atoms 1 to 6 occur in sequentially order from top to bottom. Atom positions affected by G6PD and PGI fractionation are given in blue and green, respectively. In some cases, carbon position 1 is given both as blue letter and green dot to indicate fractionation at both enzymes. Dashed arrows indicate that intermediate reactions are not shown. Wavy lines indicate fractional introduction of hydrogen from water by the PGI reaction. Note, G6PD in stem leucoplasts may additionally contribute to isotope fractionation at tree-ring glucose C-1 and H^1^. Abbreviations: F6P, fructose 6-phosphate; G6P, glucose 6-phosphate; G6PD, G6P dehydrogenase; OPPP, oxidative pentose phosphate pathway; *p*_i_, intercellular CO_2_ partial pressure; PGI, phosphoglucose isomerase; *PRE*; precipitation; RuBP, ribulose 1,5-bisphosphate.

Aside from these mechanisms, there are others that might introduce *Δ*_1_’ and *Δ*_2_’ variation. For instance, we recently reported evidence consistent with increasing relative flux through the chloroplastic OPPP in response to decreasing *p*_i_ under illumination (Fig. 5A; Wieloch *et al*., 2022a, 2023). Furthermore, under illumination, chloroplastic F6P is used for both RuBP regeneration and starch biosynthesis (Fig. 5A). Increasing *VPD* promotes photorespiration resulting in increasing RuBP regeneration relative to carbon export from the Calvin-Benson cycle into sinks such as starch.

The mechanisms described above should also introduce hydrogen isotope signals because of the hydrogen isotope effects of G6PD affecting G6P H^1^ (Hermes *et al*., 1982) and PGI affecting G6P H^2^ (Table 2, Fig. 5). However, growing season *VPD* neither correlates with *ε*_met_ pertaining to tree-ring glucose H^1^ nor H^2^ (Tables S5-6). Hence, either G6PD and PGI are not the sources of *VPD*-dependent carbon isotope fractionation in *Δ*_1_’ (and *Δ*_2_’) or the corresponding hydrogen isotope signals were washed out after introduction. Washout at H^1^ may occur during equilibration of F6P with mannose 6-phosphate by phosphomannose isomerase (*cf*. Topper, 1957). Similarly, complete washout at H^2^ may occur when the leaf-cytosolic PGI reaction is in equilibrium (Notes S3). Previously, this latter process was invoked (among others) to explain why a whole-molecule deuterium depletion observed in leaf starch was not transmitted to nocturnal sucrose (‘Introduction’; Wacker, 2022). As each conversion by PGI was found to be associated with a 0 to 50% probability for hydrogen exchange with the medium (Noltmann, 1972), partial washout of existing hydrogen isotope signals may also occur under non-equilibrium conditions (Notes S3).

In the mechanisms described above, we assumed diffusion-rubisco fractionation contributes to *VPD*-dependent *Δ_i_*’ variation. However, diffusion-rubisco fractionation affects all glucose carbon positions equally (Wieloch *et al*., 2018). Since merely two out of six glucose carbon positions carry *VPD*-dependent isotope variation, the question arises of whether the diffusion-rubisco signal was already below the detection level on introduction. If this were the case, then *VPD*-dependent *Δ*_1_’ and *Δ*_3_’ variation would originate entirely from post-rubisco processes. Furthermore, post-rubisco processes that were previously invoked to explain the absence of the diffusion-rubisco signal from C-4, C-5, and C-6 (see ‘Introduction’) would not occur.

### Isotope fractionation mechanisms in stems affecting tree-ring glucose HC-1 and HC-2

Previously, we found a change point in *ε*_met_ in 1980 (Wieloch *et al*., 2022b). Here, we found the same change point in *Δ*_1-2_’ (‘Results’). Consistent with this, *Δ*_1_’ and *Δ*_2_’ data of 1983 to 1995 exhibit a significantly lower average value and a significantly larger variance than those of 1961 to 1980 (Tables S1-2). Furthermore, *Δ*_1_’ and *Δ*_2_’ are significantly correlated during the late (Fig. 4B) but not the early period (Fig. 4A), and *ε*_met_ accounts for a significant fraction of the variance of both *Δ*_1_’ and *Δ*_2_’ during the late period (Table 4, M1, M4; Figs. 3A-B). In *Δ*_2_’, the *ε*_met_ effect is about twice as large as in *Δ*_1_’. Processes affecting *ε*_met_, *Δ*_1_’, and *Δ*_2_’ simultaneously respond to *PRE* but not *VPD* (Table S5; Table 4, M1, M2, M4, M6, M7, M11). *Δ*_1_’ and *Δ*_2_’ respond to March to July *PRE* during the late but not the early period (Table 4, M9-10). Previously, we reported evidence suggesting the groundwater table before 1980 was high enough to prevent metabolic changes causing *ε*_met_ variation (Wieloch *et al*., 2022b). By extension, this should also explain the properties of *Δ*_1_’ and *Δ*_2_’ listed above. That is, since the trees had access to groundwater during the early period, metabolic shifts that can cause intercorrelated variation in *ε*_met_, *Δ*_1_’, and *Δ*_2_’ were not induced.

Processes causing intercorrelated variation in *ε*_met_, *Δ*_1_’, and *Δ*_2_’ are probably located in the stem (see the first section of the ‘Discussion’). The *ε*_met_ signal is present at glucose H^1^ and, considerably more strongly, at H^2^ (range: 64‰ and 240‰, respectively; 1983 to 1995). In the biochemical pathway leading to tree-ring cellulose, PGI is the last enzyme acting on precursors of glucose H^2^ (Figs. 2 and 5D). With each conversion by PGI, there is a probability for hydrogen exchange with the medium of 0 to 50% (Noltmann, 1972). Thus, if we assume *Pinus nigra* stem PGI exchanges hydrogen with the medium as does spinach leaf PGI (Fedtke, 1969) and the reaction is in equilibrium, then any deuterium signal at G6P H^2^ will be washed out. Among all H-C positions of tree-ring glucose, the deuterium abundance at H^2^ is neither exceptionally high nor low during 1961 to 1980 whereas it is exceptionally high (and exceptionally variable) during 1983 to 1995 (Fig. 6). This indicates that the PGI reaction was close to or in equilibrium during 1961 to 1980 but displaced from equilibrium on the side of G6P during 1983 to 1995 (Table 2). Additionally, shifts of the PGI reaction away from equilibrium towards the side of G6P should cause ^13^C enrichment at G6P C-1 and C-2 (*Δ*_1_’ and *Δ*_2_’ decreases), and *Δ*_2_’ should decrease 3 times more than *Δ*_1_’ (Table 2). Consistent with this, we found negative relationships between *ε*_met_ and *Δ*_1_’, as well as *Δ*_2_’ (Table 4, M1 and M4). However, *Δ*_2_’ decreases only 1.88 times more than *Δ*_1_’, but this best estimate is associated with a relatively large error (SE interval: 1.04 to 3.45). That said, the offset from 3 is likely explained by increasing relative flux through the OPPP accompanying the putative PGI reaction shift (Figs. 2 and 5D). This is because G6P to 6-phosphogluconolactone conversion by G6PD exhibit ^13^C and D isotope effects 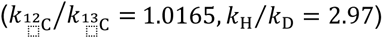 (Hermes *et al*., 1982). Hence, increasing relative OPPP flux causes ^13^C enrichment at G6P C-1 (*Δ*_1_’ decreases) and deuterium enrichment at G6P H^1^. This is consistent with both the apparently decreased PGI effects ratio (1.88 instead of 3) and, more importantly, *ε*_met_ increases at glucose H^1^ of up to 64‰ during 1983 to 1995 (Fig. 6).

**Figure 6.**
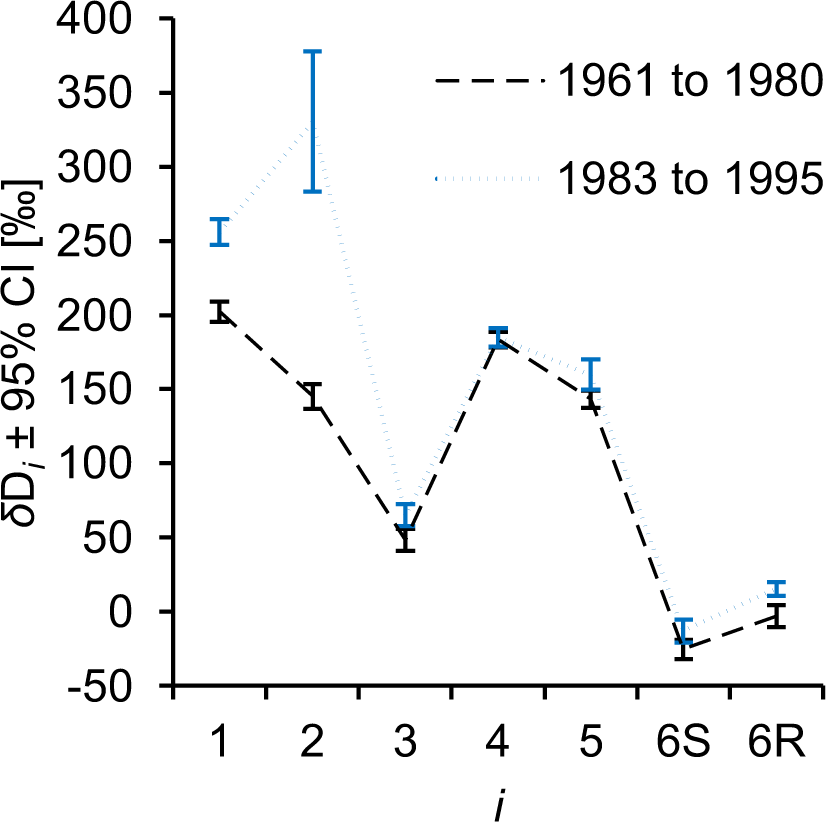
Average intramolecular *δ*D*_i_* patterns of the periods 1961 to 1980 and 1983 to 1995 (black and blue, respectively). The data were acquired for tree-ring glucose of *Pinus nigra* laid down at a site in the Vienna basin. The figure shows discrete data. Dashed and dotted lines were added to guide the eye. Data reference: Average deuterium abundance of the methyl-group hydrogens of the glucose derivative used for NMRS measurements. Modified figure from Wieloch *et al*. (2022b).

Sucrose translocated from leaves can be split into UDP-glucose and fructose via sucrose synthase or glucose and fructose via invertase (Fig. 2). UDP-glucose entering tree-ring cellulose biosynthesis directly via sucrose synthase is protected from isotope fractionation by PGI and G6PD. However, in stems of juvenile *Quercus petraea* and *Picea abies*, at least 79% and 43% of the precursors of tree-ring glucose went through PGI catalysis, respectively (Augusti *et al*., 2006). Theoretically, shifts of the PGI reaction away from equilibrium towards the side of G6P can cause *ε*_met_ increases at glucose H^2^ of up to 611‰ (Notes S3, hydrogen exchange with the medium not considered). With 43% and 79% of all precursors of tree-ring glucose undergoing PGI catalysis, *ε*_met_ increases at glucose H^2^ of up to 263‰ and 483‰ are possible, respectively. Thus, the PGI-related fractionation mechanism proposed here can potentially cause previously reported *ε*_met_ increases at glucose H^2^ of up to 240‰ (Wieloch *et al*., 2022b). Shifts in sucrose cleavage by sucrose synthase versus invertase may exert additional control on the *ε*_met_ signal at glucose H^2^.

Based on results and interpretations presented above, decreasing stem water content is associated with both increasing OPPP flux and a shift of the PGI reaction away from equilibrium towards the side of G6P corresponding to low relative F6P concentration (Fig. 5D). We propose these concerted shifts may ensure redox homeostasis and balanced substrate supply to glycolysis as follows. In heterotrophic tissue, NADPH from the OPPP is believed to be central for maintaining redox homeostasis (Fig. 2) (Stincone *et al*., 2015). Flux through the OPPP is regulated at G6PD. Heterotrophic G6PD activity reportedly increases with drought (Liu *et al*., 2013; Wang *et al*., 2016, 2020), oxidative load (Wang *et al*., 2016, 2020; Li *et al*., 2020), NADPH demand (Wendt *et al*., 2000; Esposito *et al*., 2001; Castiglia *et al*., 2015), and abscisic acid concentration (Cardi *et al*., 2011; Wang *et al*., 2016). Decreasing stem water content may cause increasing OPPP flux via increasing abscisic acid concentration (Brunetti *et al*., 2020), and possibly increasing oxidative load increasing the demand for NADPH. In turn, increasing OPPP flux results in increasing supply of pentose phosphates which may feed into glycolysis via the reductive part of the pentose phosphate pathway (Figs. 2 and 5D). This would reduce the demand for glycolytic substrates supplied via PGI. The shift of the PGI reaction away from equilibrium towards the side of G6P may reflect this decreased demand and result from PGI downregulation by intermediates of the pentose phosphate pathway such as erythrose 4-phosphate, ribulose 5-phosphate, and 6-phosphogluconate (Parr, 1956; Grazi *et al*., 1960; Salas *et al*., 1965). Furthermore, relative changes in G6P-to-F6P supply versus consumption may contribute to the shift of the PGI reaction. For instance, while starch storage consumes G6P, remobilisation supplies G6P (Noronha *et al*., 2018). Under drought, the storage-to-remobilisation balance may tilt towards remobilisation (Mitchell *et al*., 2013; Thalmann & Santelia, 2017; Tsamir-Rimon *et al*., 2021). Consequently, the PGI reaction may move towards the side of G6P. Similarly, we previously reported below-average tree-ring widths for years in which the PGI reaction is on the side of G6P (Wieloch *et al*., 2022b). Hence, in these years, G6P consumption by growth may have been reduced while F6P consumption by downstream metabolism may have been maintained.

### 1961 to 1980: A period of homeostasis with respect to processes affecting Δ_1_’, Δ_2_’, Δ_3_’, and ε_met_

During 1961 to 1980, *Δ*_1_’, *Δ*_2_’, and *Δ*_3_’ are not significantly correlated which contrasts with the period 1983 to 1995 (Figs. 4a-b). Similarly, relationships of *Δ*_1_’ and *Δ*_3_’ with *VPD* and *Δ*_1_’ and *Δ*_2_’ with *PRE* observed during the late period are largely absent during the early period (Tables 3-5) even though there is no difference in the magnitude of *VPD* and *PRE* variability between these periods (Fig. S1). Like *Δ*_1-2_’ and *Δ*_1-3_’, *ε*_met_ exhibits a change point in 1980 and responds to *PRE* after but not before 1980 (Wieloch *et al*., 2022b). This shift in *ε*_met_ sensitivity was attributed to long-term drought which intensified over the study period and proposedly lead to a groundwater depletion below a critical level in 1980 (Wieloch *et al*., 2022b). By extension, this groundwater depletion might also explain the insensitivity of *Δ*_1_’ and *Δ*_3_’ to *VPD* and *Δ*_1_’ and *Δ*_2_’ to *PRE* during 1961 to 1980 and their sensitivity from 1983 onwards. Thus, while the trees had access to groundwater, leaf- and stem-level processes affecting *Δ*_1_’, *Δ*_2_’, *Δ*_3_’ and *ε*_met_ could apparently maintain homeostasis despite changing atmospheric conditions.

### Isotope fractionation mechanisms in leaves affecting tree-ring glucose C-5 and C-6

No change points were detected in *Δ*_5_’ and *Δ*_6_’ (‘Results’; Tables S1-2). Furthermore, *Δ*_5_’ and *Δ*_6_’ remain significantly correlated across the entire study period (Figs. 1B and 4A-B), and *RAD* is the most influential environmental cofactor (Table 6). Models including *RAD* and *TMP* as cofactors capture most of the systematic variance in *Δ*_5-6_’, *Δ*_5_’, and *Δ*_6_’ (Table 7, M1-3; Table S4). These relationships hold for both the early and late study period (Table 7, M4-5) with *Δ*_5_’ effects being about 1.5-fold larger than *Δ*_6_’ effects (M2 versus M3, SE interval: 1.1 to 2.28).

Previously, we reported a negative relationship between tree-ring glucose *Δ*_5-6_’ and reconstructed tropospheric O_3_ concentration (‘Introduction’, Wieloch *et al*., 2022c). Light stimulates tropospheric O_3_ formation (Lu *et al*., 2019). This may explain the negative relationship between tree-ring glucose *Δ*_5-6_’ and *RAD* reported here (Table 7, M1-3). Furthermore, we previously explained the absence of the diffusion-rubisco signal from glucose C-5 and C-6 (*inter alia*) by interaction between photorespiration and the TCAC (‘Introduction’, Wieloch *et al*., 2022c). As *TMP* increases, photorespiration increases more than photosynthesis (Long, 1991). This may result in decreasing flux of PEP into the TCAC (‘Introduction’) and explain the positive relationship between tree-ring glucose *Δ*_5-6_’ and *TMP* reported here (Table 7, M1-3).

### Isotope fractionation mechanisms in leaves affecting tree-ring glucose C-4

As for *Δ*_5_’ and *Δ*_6_’, no change point was detected in *Δ*_4_’ (‘Results’; Tables S1-2). Considering the entire study period, *Δ*_4_’ is weakly associated with *Δ*_5-6_’ (Fig. 1B). Consistent with this, the *Δ*_5-6_’-climate model works reasonably well for *Δ*_4_’ considering the relatively low systematic variance in *Δ*_4_’ of 38% (Table 7, M1 and M6; Table S4). Introduction of the *Δ*_4_’ and *Δ*_5-6_’ signals proposedly involves leaf-level consumption of PGA and PEP by downstream metabolism, respectively (Wieloch *et al*., 2021, 2022c). Since PGA is a precursor of PEP (Fig. 2), our previously proposed theories of signal introduction are in line with the observation that *Δ*_4_’, *Δ*_5_’ and *Δ*_6_’ are associated and respond to the same environmental parameters.

## Conclusions and future directions

Dual-isotope-environment analysis was used to deconvolute isotope signals and provide several new insights into plant isotope fractionation. First, the diffusion-rubisco signal was previously shown to be absent from tree-ring glucose C-4 to C-6 (Wieloch *et al*., 2021, 2022c) but believed to be present at C-1 to C-3 (Wieloch *et al*., 2018). Here, this signal was found to also be absent from C-2. Second, isotope fractionation beyond leaves is commonly considered to be constant for any given species (Roden *et al*., 2000; Gagen *et al*., 2022). However, our results suggest a significant part of the carbon and hydrogen isotope variation in tree-ring glucose originates in stems from processes affecting *Δ*_1_’, *Δ*_2_’, and *ε*_met_ simultaneously. Third, *VPD* affects *Δ*_1_’ and *Δ*_3_’ and *PRE* affects *Δ*_1_’, *Δ*_2_’, and *ε*_met_ (Table 4). These relationships proposedly reflect water content variability in leaves and stems, respectively. They apply to the late but not the early study period consistent with the finding of a change point in both the *ε*_met_ (Wieloch *et al*., 2022b) and *Δ*_1-3_’ series (see above). This change point proposedly marks the crossing of a physiologically relevant groundwater threshold (Wieloch *et al*., 2022b). Additionally, we reported *Δ*_5-6_’ relationships with *RAD* and *TMP* which apply to the entire study period (Table 4). These latter relationships are consistent with previously proposed isotope fractionation mechanisms (Wieloch *et al*., 2022c). By contrast, we here revised and expanded our previous theory on the mechanisms introducing *Δ*_1_’, *Δ*_2_’, *Δ*_3_’, and *ε*_met_ variability. Given the multitude of isotope-environment relationships (including change-point responses), intramolecular carbon isotope analysis has a remarkable potential for reconstructions of environmental conditions (*VPD*, *PRE*, *RAD*, *TMP*, soil moisture, groundwater thresholds, tropospheric O_3_ concentration), tissue water content (leaf, stem), metabolic flux variability (various processes), and ecophysiological properties such as intrinsic water-use efficiency across space and time. Complementing hydrogen isotope analysis is expected to significantly enhance these capabilities.

Understanding isotope fractionation mechanisms is central for retrospective studies of plant physiology and climate based on tree-ring isotope data, and there is considerable room for improvement as shown above. Research in several largely unexploited areas is needed to make progress. First, there is a basic need for more *in vitro* data on intramolecular isotope effects of enzyme reactions including the reactions catalysed by triosephosphate isomerase, transketolase, PEPC, PK, and DAHPS. Second, intramolecular isotope analyses of leaf metabolites including starch and sucrose from both controlled and natural environments are needed to generate a baseline for mechanistic studies of isotope fractionation along the pathway from leaves to wood. Additionally, intramolecular isotope analysis of metabolites from wood slices acclimated to different ambient conditions (e.g., wet versus dry, varying sucrose supply) will be insightful. Third, combined analysis of intramolecular ^13^C and deuterium data can help to separate isotope signals. Fourth, genetic modification of key enzymes may help to test proposed isotope fractionation mechanisms *in vivo*. Fifth, intramolecular isotope fractionation affecting tree-ring glucose is complex (see above). Software programs such as QIRN enable the convenient simulation of natural isotope abundances in complex metabolic networks (Mueller *et al*., 2022). If expanded, these programs may help to extract metabolic information from intramolecular tree-ring isotope data. This would require routines enabling control of relative flux at metabolic branchpoints by optimising regressions between (i) relative branchpoint flux and environmental parameters and (ii) simulated and observed isotope data. In summary, intramolecular isotope analysis has an enormous potential to advance our knowledge about isotope fractionation mechanisms, plant ecophysiology, and paleoclimatology.

## Acknowledgements

TW’s work was carried out with funding from “Formas - a Swedish Research Council for Sustainable Development” (2022-02833, Grant recipient: TW). MHP’s work was supported by the Swiss National Science Foundation (205492).

## Author Contributions

Conceptualisation: TW; Investigation: TW with input from all authors; Visualization: TW; Development of isotope theory: TW; Project administration: TW; Writing: TW with input from all authors.

## Competing interests

None declared.

## Data availability

The authors declare that the data supporting the findings of this study are available within the paper and its supplementary information files.

## Notes

### Competing Interest Statement

The authors have declared no competing interest.

### Summary of Updates

- major revisions of main text and supporting information - Figure 5 added

