## Supporting Information for "New insights into the mechanisms of plant isotope fractionation from combined analysis of intramolecular ^13^C and deuterium abundances in *Pinus nigra* tree-ring glucose"

The following Supporting Information is available for this article:

**Notes S1.** Materials and Methods (expanded).

**Notes S2.** Hydro-carbon isotope fractionation during 1961 to 1980.

**Notes S3.** Estimated deuterium fractionation due to shifts of the phosphoglucose isomerase reaction.

**Table S1.** Shapiro-Wilk normality test.

**Table S2.** F and T test.

**Table S3.** Pearson correlations between $\Delta_i$' and $\varepsilon_{met}$ series of the period 1983 to 1995.

**Table S4.** Components of variance in $\Delta_i$' series.

**Table S5.** Pearson correlation coefficients and associated levels of significance of $\varepsilon_{met}$-climate relationships for the period 1983 to 1995.

**Table S6.** Linear regression model of $\varepsilon_{met\ (H1)}$ as function of growing season air vapour pressure deficit and March to July precipitation.

**Figure S1.** Air vapour pressure deficit of the growing season and March to July precipitation over the period 1961 to 1995 in the Vienna basin.

**Notes S1. Materials and Methods (expanded).**

The tree-ring samples used here were described previously (Leal *et al.*, 2008). Two tree-ring cores were taken from each of 19 *Pinus nigra* Arnold trees at Bierhäuselberg (Vienna region, Austria, 48.13° N, 16.23° E, 350 m AMSL). The site is unmanaged, has an open canopy, and shallow soil. Sampling targeted dominant trees with umbrella-shaped crowns indicating frequent water deficits. Tree rings were dated by standard dendrochronological methods (Speer, 2010). Generation of the $\Delta_i$' and $\varepsilon_{met}$ datasets was described previously (Wieloch *et al.*, 2018, 2022). On average, tree-ring width series started in 1865 (range: 1840 to 1918). To preclude significant isotope shifts due to developmental processes or the contribution of soil-respired $CO_2$, our isotope analyses start in 1961. At this point, all trees had reached a stable position in the canopy. Since all tree-ring material of each year was combined into annual pools before measuring isotopes, our data represent the species at the site rather than individual trees. To ensure high precision, glucose derivative samples < 20 mg were excluded from measurement (1977, 1978, 1981, and 1982).

Annual $p_a$ data were obtained for the Mauna Loa Observatory, Hawaii (curators: Pieter Tans, NOAA/ESRL, Boulder, USA; Ralph Keeling, Scripps Institution of Oceanography, La Jolla, USA). $\Delta$ may be affected by $p_a$ (Schubert & Jahren, 2012). However, in the present case, $\Delta$ and $p_a$ are not significantly correlated ($r$ = -0.32, $p$ > 0.05, $n$ = 31). Moreover, in line with the mechanistic model of diffusion-rubisco $^{13}C$ discrimination (Farquhar *et al.*, 1989), most previous studies have reported positive slopes for the regression of these variables (Schubert & Jahren, 2012), but we found a negative slope (-0.02 ± 0.01SE). Thus, the effect of $p_a$ on $\Delta$ is not verifiable here, and we did not correct our data for $p_a$.

By contrast, $\Delta_i$' was corrected for $^{13}C$ signal redistribution by heterotrophic triose phosphate cycling (Wieloch *et al.*, 2018). In this process, part of the sucrose molecules entering tree-ring cells are broken down into the triose phosphates glyceraldehyde 3-phosphate and dihydroxyacetone phosphate. These molecules are in chemical equilibrium which causes $^{13}C$ signal transfer among carbon positions. Consequently, tree-ring glucose synthesised from triose phosphates inherits a rearranged intramolecular $^{13}C$ signal distribution. Correcting $\Delta_i$' for heterotrophic triose phosphate cycling restores the original $^{13}C$ signal distribution.

Estimation of $\varepsilon_{met}$ aims to remove fractionations by non-metabolic processes such as leaf water deuterium enrichment (Wieloch *et al.*, 2022). Furthermore, $\varepsilon_{met}$ series calculated separately for $H^1$ and $H^2$ (by replacing the numerator in equation 1 with $D_1$ and $D_2$, respectively) are essentially perfectly correlated considering that both series exhibit significant random variation due to the relatively large error of NMRS measurements (1983 to 1995, $r$ = 0.92, $p$ = $10^{-5}$, $n$ = 13).

Based on *TMP* during the study period (1961 to 1995), the growing season at the site was estimated to extend from March to November (Wieloch *et al.*, 2018). Conifers form tree rings over the course of several months (Cuny *et al.*, 2014). Therefore, all statistical analyses exclusively consider periods comprising ≥ 4 growing season months. According to autocorrelation analyses on $\Delta_i$' and $\Delta$ series, the growth of the trees studied here has not been significantly affect by interannual carry-over of carbon (Wieloch *et al.*, 2018). Therefore, our statistical analyses do not consider the climate conditions of previous years.

After mean-centring and unit-variance scaling of $\Delta_i$' series, hierarchical cluster analysis was done with the functions dist() and hclust() of the 'stats' package in R (R Core Team, 2021) choosing Euclidean distances and Ward's fusion criterion as inputs (Ward, 1963). Pearson correlation analysis, ordinary least squares regression analysis, and Shapiro-Wilk normality tests were respectively done with the functions cor(), lm() and shapiro.test() of the 'stats' package in R (R Core Team, 2021). The fraction of systematic variance in isotope series was estimated according to published procedures (Nilsson *et al.*, 1996). Change point tests were done with the function detectChangePointBatch() of the 'cpm' package in R (parametric generalised likelihood ratio test, nonparametric Lepage test) (Ross, 2015). F tests and one-tailed T tests were respectively done with the functions f.test() and t.test() in Excel (Microsoft Corporation, Redmond, WA, USA).

**Notes S2. Hydro-carbon isotope fractionation during 1961 to 1980.**

This note investigates whether, during the period 1961 to 1980, $\Delta_1$', $\Delta_2$', and $\Delta_3$' contain systematic variation and whether there is correlated hydrogen and carbon (hydro-carbon) isotope fractionation.

The variance of $\Delta_1$' and $\Delta_2$' during 1961 to 1980 is low compared to 1983 to 1995 ($\Delta_1$': 1.90‰ versus 5.86‰; $\Delta_2$': 1.25‰ versus 5.84‰) and only slightly exceeds the estimated random error variance ($\Delta_1$': 1.12‰; $\Delta_2$': 0.9‰). By contrast, $\Delta_3$' exhibits higher variance during 1961 to 1980 than during 1983 to 1995 (2.43‰ versus 1.34‰) and exceeds the estimated random error variance by about a factor of two (1.06‰). In relation to the total variance, $\Delta_1$', $\Delta_2$', and $\Delta_3$' exhibit 41%, 28%, and 56% systematic variance during 1961 to 1980, respectively.

Furthermore, during 1961 to 1980, $\Delta_1$', $\Delta_2$', and $\Delta_3$' are not significantly correlated (Fig. 4B, Table N1). Similarly, metabolic hydrogen isotope fractionation at glucose $H^2$, $\varepsilon_{met\ (H2)}$, is neither significantly correlated with $\Delta_1$', $\Delta_2$', and $\Delta_3$' nor with metabolic hydrogen isotope fractionation at glucose $H^1$, $\varepsilon_{met\ (H1)}$. By contrast, we found indications for significant negative correlation between $\Delta_1$' and $\varepsilon_{met\ (H1)}$ (Table N1, $r$ = -0.68, $p$ < 0.01, $n$ = 18). However, looking at the corresponding scatterplot revealed an outlier (Fig. N1, red circle), and removing this outlier from the data removes the correlation between $\Delta_1$' and $\varepsilon_{met\ (H1)}$ ($r$ = -0.39, $p$ > 0.01, $n$ = 17). Thus, there is no evidence for hydro-carbon isotope fractionation during 1961 to 1980.

**Table N1 Correlation among isotope series during 1961 to 1980 (*n* = 18, missing years: 1977, 1978)**

| | $\Delta_1$' | $\Delta_2$' | $\Delta_3$' | $\varepsilon_{met\ (H1)}$ | $\varepsilon_{met\ (H2)}$ |
|---|---|---|---|---|---|
| $\Delta_1$' | 1.00 | | | | |
| $\Delta_2$' | -0.21 | 1.00 | | | |
| $\Delta_3$' | -0.25 | 0.41 | 1.00 | | |
| $\varepsilon_{met\ (H1)}$ | -0.68** | 0.12 | 0.19 | 1.00 | |
| $\varepsilon_{met\ (H2)}$ | 0.01 | 0.24 | 0.09 | 0.05 | 1.00 |

$\varepsilon_{met\ (H1)}$, $\varepsilon_{met\ (H2)}$, $\Delta_1$', $\Delta_2$', and $\Delta_3$' denote hydrogen isotope fractionation caused by metabolic processes at glucose $H^1$ and $H^2$, and $^{13}C$ discrimination at glucose C-1, C-2, and C-3, respectively. Glucose was extracted across an annually resolved tree-ring series of *Pinus nigra* from the Vienna Basin. Significance of series correlation: **, $p \leq 0.01$.

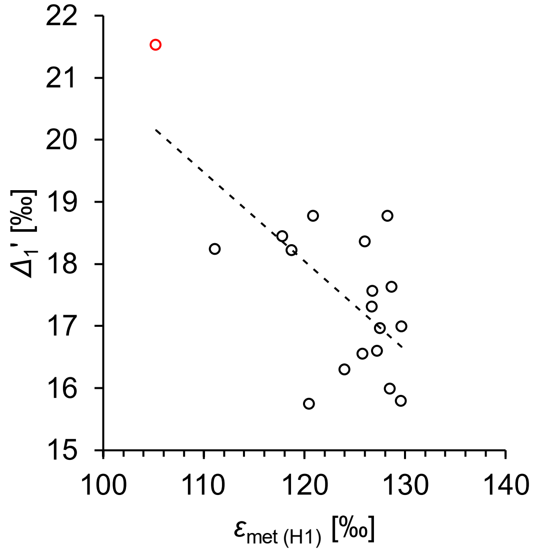


**Figure N1. Relationship between metabolic hydrogen isotope fractionation at tree-ring glucose $H^1$ ($\varepsilon_{met\ (H1)}$) and $^{13}C$ discrimination at C-1 ($\Delta_1$') during 1961 to 1980.** Dashed line, relationship between $\varepsilon_{met\ (H1)}$ and $\Delta_1$'. Red circle, outlier.

**Notes S3. Estimated deuterium fractionation due to shifts of the phosphoglucose isomerase reaction.**

Table 2 shows reported hydrogen isotope effects of the reaction catalysed by phosphoglucose isomerase (PGI). Full expression of the kinetic isotope effect in the direction glucose 6-phosphate (G6P) to fructose 6-phosphate (F6P) results in 500‰ relative deuterium depletion in F6P as

$$\varepsilon_{\mathrm{inv}} = \frac{1}{\alpha} - 1 = \frac{k_{\mathrm{D}}}{k_{\mathrm{H}}} - 1 = \frac{k_{\mathrm{D}} - k_{\mathrm{H}}}{k_{\mathrm{H}}} = \frac{1}{2} - 1 = -0.5 \qquad \text{(S1)}$$

where $\varepsilon_{\mathrm{inv}}$ denotes deuterium fractionation, $\alpha$ denotes hydrogen isotope effect, and $k_{\mathrm{D}}$ and $k_{\mathrm{H}}$ denote reaction rate of the heavy and light G6P isotopologue, respectively. By contrast, full expression of the equilibrium isotope effect results in 111‰ relative deuterium enrichment in F6P as

$$\varepsilon_{\mathrm{inv}} = \frac{1}{\alpha} - 1 = \frac{1}{0.9} - 1 = 0.\bar{1} \qquad \text{(S2)}$$

Hence, in the absence of hydrogen exchange with the medium, shifts of the PGI reaction away from equilibrium towards the side of G6P can cause 611‰ deuterium depletion in F6P and 611‰ deuterium enrichment in G6P.

That said, each conversion by PGI was found to be associated with a 0 to 50% probability for hydrogen exchange with the medium (see Fig. 5; Noltmann, 1972). For instance, in spinach leaf extracts, hydrogen transfer in the F6P-to-G6P direction reportedly proceeds with ≈30% of the hydrogen exchanging with the medium, i.e., ≈70% are transferred intramolecularly (Fedtke, 1969). In chlorella extracts, on the other hand, hydrogen transfer occurs 100% intramolecularly, i.e., without exchange with the medium (Simon *et al.*, 1964; Dorrer *et al.*, 1966; Fedtke, 1969). Full intramolecularity may proposedly result if PGI is embedded in a metabolon, a transient multi-enzyme complex (Simon *et al.*, 1964). Additionally, temperature was found to affect the degree of hydrogen exchange with the medium (Rose & O’Connell, 1961; Fedtke, 1969). Taken together, deuterium fractionation by PGI might be dampened by fractional hydrogen incorporation from the medium.

**Table S1. Shapiro-Wilk normality test (*n* = 31).**

| | $\Delta_1'$ | $\Delta_2'$ | $\Delta_3'$ | $\Delta_4'$ | $\Delta_5'$ | $\Delta_6'$ | $\Delta$ | $\Delta_{1\text{-}2}'$ | $\Delta_{1\text{-}3}'$ | $\Delta_{5\text{-}6}'$ | $\Delta_{4\text{-}6}'$ |
|---|---|---|---|---|---|---|---|---|---|---|---|
| *W* | 0.93 | 0.97 | 0.99 | 0.95 | 0.96 | 0.94 | 0.91 | 0.90 | 0.90 | 0.97 | 0.96 |
| *p* | **0.04** | 0.47 | 1.00 | 0.13 | 0.23 | 0.06 | **0.01** | **0.01** | **0.01** | 0.64 | 0.35 |

**Table S2. F and T test (*n* = 18 and 13).**

| | $\Delta_1'$ | $\Delta_2'$ | $\Delta_3'$ | $\Delta_4'$ | $\Delta_5'$ | $\Delta_6'$ | $\Delta$ | $\Delta_{1\text{-}2}'$ | $\Delta_{1\text{-}3}'$ | $\Delta_{5\text{-}6}'$ | $\Delta_{4\text{-}6}'$ |
|---|---|---|---|---|---|---|---|---|---|---|---|
| **Variance, 1961-1980** | 1.90 | 1.25 | 2.43 | 1.38 | 3.18 | 1.86 | 0.30 | 0.63 | 0.60 | 2.02 | 1.16 |
| **Variance, 1983-1995** | 5.86 | 5.84 | 1.34 | 0.99 | 2.44 | 1.89 | 1.47 | 4.89 | 3.25 | 1.86 | 1.21 |
| **F test, *p* =** | **0.03** | **0.004** | 0.32 | 0.59 | 0.68 | 0.92 | **0.003** | **0.001** | **0.002** | 0.94 | 0.88 |
| **Average, 1961-1980** | 17.5 | 7.8 | 6.4 | 13.5 | 19.9 | 20.4 | 14.3 | 12.6 | 10.5 | 20.2 | 17.9 |
| **Average, 1983-1995** | 15.8 | 6.0 | 6.0 | 14.2 | 20.1 | 19.6 | 13.6 | 10.8 | 9.2 | 19.9 | 18.0 |
| **one-tailed T test, *p* =** | **0.02** | **0.01** | 0.20 | 0.06 | 0.37 | 0.06 | **0.05** | **0.01** | **0.01** | 0.28 | 0.48 |

**Table S3. Pearson correlations between $\Delta_i$' and $\varepsilon_{met}$ series of the period 1983 to 1995 (*n* = 13).**

| | $\Delta_1'$ | $\Delta_2'$ | $\Delta_3'$ | $\Delta_4'$ | $\Delta_5'$ | $\Delta_6'$ |
|---|---|---|---|---|---|---|
| *r* | -0.59 | -0.74 | -0.45 | -0.05 | -0.05 | 0.07 |
| *p* | 0.03 | 0.004 | 0.12 | 0.87 | 0.88 | 0.81 |

**Table S4. Components of variance in $\Delta_i$' series.**

| | 1983 to 1995 | | | 1964 to 1995 | | | |
|---|---|---|---|---|---|---|---|
| | $\Delta_1'$ | $\Delta_2'$ | $\Delta_3'$ | $\Delta_4'$ | $\Delta_5'$ | $\Delta_6'$ | $\Delta_{5\text{-}6}'$ |
| **total variance (‰)** | 5.86 | 5.84 | 1.34 | 1.37 | 2.86 | 1.76 | 1.80 |
| **random variance (‰)** | 0.71 | 0.73 | 0.71 | 0.85 | 0.96 | 0.97 | 0.48 |
| **fraction of systematic variance (%)** | 88 | 88 | 47 | 38 | 66 | 45 | 73 |

Random variance due to measurement errors in $\Delta_i$' was estimated according to published procedures (Nilsson *et al.*, 1996).

**Table S5. Pearson correlation coefficients and associated levels of significance of $\varepsilon_{met}$-climate relationships for the period 1983 to 1995 ($n$ = 13).**

| | *VPD* | *PRE* | $SPEI_1$ | $SPEI_3$ | $SPEI_4$ | $SPEI_6$ | $SPEI_8$ | $SPEI_{12}$ | $SPEI_{16}$ | $SPEI_{24}$ | $SPEI_{36}$ | $SPEI_{48}$ | *TMP* | *SD* | *RAD* |
|---|---|---|---|---|---|---|---|---|---|---|---|---|---|---|---|
| **MAMJ** | 0.55 | -0.70 | -0.37 | -0.17 | -0.13 | -0.16 | -0.20 | 0.09 | 0.22 | 0.07 | 0.04 | 0.20 | 0.15 | -0.07 | -0.22 |
| **MAMJJ** | 0.42 | -0.84 | -0.43 | -0.26 | -0.24 | -0.24 | -0.26 | 0.00 | 0.16 | 0.04 | 0.00 | 0.15 | -0.04 | -0.22 | -0.37 |
| **MAMJJA** | 0.29 | -0.66 | -0.27 | -0.26 | -0.27 | -0.27 | -0.26 | -0.08 | 0.14 | 0.04 | -0.02 | 0.11 | -0.03 | -0.12 | -0.31 |
| **MAMJJAS** | 0.24 | -0.60 | -0.20 | -0.22 | -0.25 | -0.29 | -0.26 | -0.14 | 0.10 | 0.03 | -0.02 | 0.08 | -0.13 | -0.10 | -0.31 |
| **MAMJJASO** | 0.28 | -0.63 | -0.24 | -0.17 | -0.23 | -0.29 | -0.28 | -0.16 | 0.07 | 0.02 | -0.03 | 0.06 | -0.09 | -0.05 | -0.31 |
| **MAMJJASON** | 0.27 | -0.67 | -0.27 | -0.17 | -0.20 | -0.29 | -0.29 | -0.18 | 0.03 | 0.02 | -0.04 | 0.03 | -0.14 | 0.01 | -0.33 |
| **AMJJ** | 0.41 | -0.70 | -0.41 | -0.37 | -0.32 | -0.26 | -0.36 | -0.09 | 0.10 | 0.01 | -0.03 | 0.12 | 0.03 | -0.17 | -0.36 |
| **AMJJA** | 0.28 | -0.50 | -0.28 | -0.36 | -0.32 | -0.28 | -0.33 | -0.17 | 0.08 | 0.01 | -0.04 | 0.08 | 0.02 | -0.07 | -0.31 |
| **AMJJAS** | 0.23 | -0.47 | -0.20 | -0.31 | -0.30 | -0.29 | -0.31 | -0.22 | 0.05 | 0.01 | -0.04 | 0.05 | -0.11 | -0.05 | -0.29 |
| **AMJJASO** | 0.27 | -0.51 | -0.24 | -0.25 | -0.28 | -0.29 | -0.32 | -0.23 | 0.01 | 0.00 | -0.05 | 0.03 | -0.05 | 0.00 | -0.29 |
| **AMJJASON** | 0.26 | -0.59 | -0.28 | -0.24 | -0.23 | -0.29 | -0.32 | -0.24 | -0.02 | 0.01 | -0.06 | 0.00 | -0.13 | 0.07 | -0.30 |
| **MJJA** | 0.22 | -0.42 | -0.23 | -0.40 | -0.37 | -0.30 | -0.33 | -0.28 | 0.01 | -0.02 | -0.07 | 0.06 | -0.03 | -0.06 | -0.29 |
| **MJJAS** | 0.17 | -0.43 | -0.17 | -0.34 | -0.34 | -0.30 | -0.30 | -0.30 | -0.02 | -0.03 | -0.07 | 0.03 | -0.17 | -0.03 | -0.28 |
| **MJJASO** | 0.21 | -0.46 | -0.21 | -0.28 | -0.31 | -0.30 | -0.31 | -0.29 | -0.06 | -0.03 | -0.07 | 0.01 | -0.11 | 0.02 | -0.28 |
| **MJJASON** | 0.20 | -0.53 | -0.24 | -0.27 | -0.26 | -0.29 | -0.32 | -0.29 | -0.08 | -0.02 | -0.07 | -0.02 | -0.19 | 0.10 | -0.29 |
| **JJAS** | 0.08 | -0.17 | -0.08 | -0.31 | -0.36 | -0.31 | -0.28 | -0.32 | -0.06 | -0.02 | -0.09 | -0.01 | -0.41 | -0.08 | -0.34 |
| **JJASO** | 0.13 | -0.16 | -0.09 | -0.23 | -0.31 | -0.31 | -0.29 | -0.31 | -0.10 | -0.03 | -0.09 | -0.03 | -0.40 | -0.01 | -0.33 |
| **JJASON** | 0.13 | -0.25 | -0.14 | -0.22 | -0.26 | -0.29 | -0.29 | -0.30 | -0.12 | -0.01 | -0.09 | -0.05 | -0.40 | 0.09 | -0.35 |
| **JASO** | 0.06 | 0.01 | 0.07 | -0.13 | -0.26 | -0.31 | -0.27 | -0.30 | -0.13 | -0.03 | -0.09 | -0.07 | -0.40 | -0.02 | -0.32 |
| **JASON** | 0.06 | -0.09 | -0.02 | -0.11 | -0.19 | -0.29 | -0.28 | -0.29 | -0.15 | -0.01 | -0.09 | -0.09 | -0.41 | 0.09 | -0.33 |
| **ASON** | 0.08 | 0.10 | 0.14 | 0.03 | -0.09 | -0.27 | -0.25 | -0.27 | -0.15 | 0.00 | -0.08 | -0.10 | -0.24 | 0.38 | -0.12 |

Significance levels: ≤ 0.05, light grey; ≤ 0.01, grey; ≤ 0.001, dark grey. Climate parameters: *PRE*, precipitation; *RAD*, global radiation; *SD*, sunshine duration; $SPEI_i$, standardised precipitation-evapotranspiration index of different periods ($i$ = 1, 3, 6, 8, 12, 16, 24, 36, 48 months); *TMP*, air temperature; *VPD*, air vapour pressure deficit. Climate data were averaged for all ≥ 4-month periods of the growing season (March to November). Months were abbreviated by their initial letters. $\varepsilon_{met}$ denotes hydrogen isotope fractionation caused by metabolic processes at glucose $H^1$ and $H^2$. Glucose was extracted across an annually resolved tree-ring series of *Pinus nigra*. Data of the period March to November have been published previously (Wieloch *et al.*, 2022).

**Table S6. Linear regression model of $\varepsilon_{met\ (H1)}$ and $\varepsilon_{met\ (H2)}$ as function of March to July precipitation (*PRE*) and growing season air vapour pressure deficit (*VPD*).**

| $\varepsilon_{met\ (H1)}$ ~ ***PRE* + *VPD*, 1983-1995** | | | |
|---|---|---|---|
| $R^2 = 0.6$, $adjR^2 = 0.52$, $p = 0.01$, $n = 13$ | | | |
| | **Estimate** | **± SE** | ***p* ≤** |
| **Intercept** | 209 | 53 | 0.003 |
| ***PRE*** | -0.248 | 0.074 | 0.007 |
| ***VPD*** | 0.0221 | 0.0694 | 0.76 |
| $\varepsilon_{met\ (H2)}$ ~ ***PRE* + *VPD*, 1983-1995** | | | |
| $R^2 = 0.73$, $adjR^2 = 0.67$, $p < 0.002$, $n = 13$ | | | |
| | **Estimate** | **± SE** | ***p* ≤** |
| **Intercept** | 665 | 190 | 0.006 |
| ***PRE*** | -1.31 | 0.27 | 0.001 |
| ***VPD*** | -0.186 | 0.250 | 0.47 |

$\varepsilon_{met\ (H1)}$ and $\varepsilon_{met\ (H2)}$ denote hydrogen isotope fractionation caused by metabolic processes at glucose $H^1$ and $H^2$, respectively. Glucose was extracted across an annually resolved tree-ring series of *Pinus nigra* from the Vienna Basin.

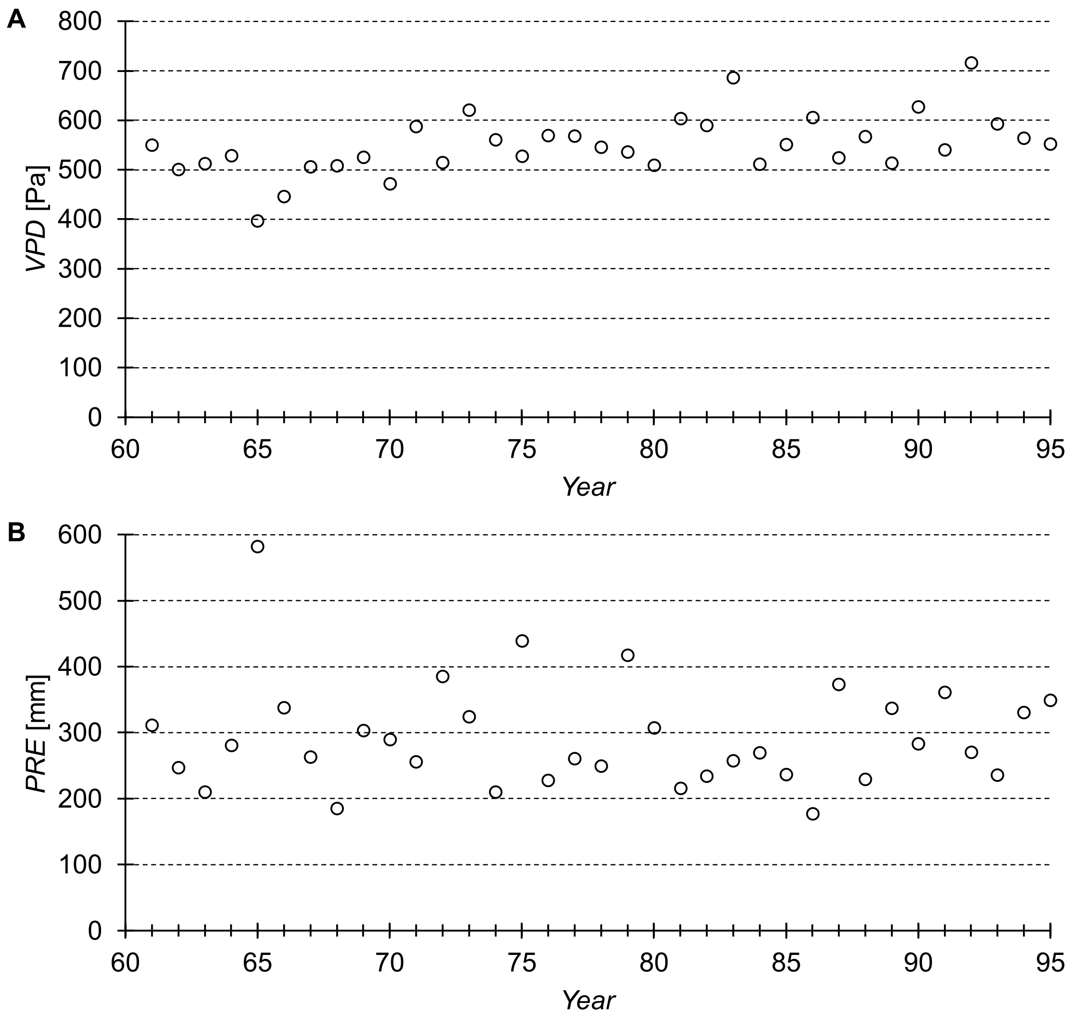


**Figure S1. Air vapour pressure deficit (*VPD*) of the growing season (A) and March to July precipitation (B, *PRE*) over the period 1961 to 1995 in the Vienna basin.** Raw data were measured at the climate station Hohe Warte (Central Institution for Meteorology and Geodynamics, Vienna, Austria, 48.23° N, 16.35° E, 198 m AMSL, WMO ID: 1103500) (Klein Tank *et al.*, 2002). *VPD* was calculated following published procedures (Abtew & Melesse, 2013).
